# Sustained fertility from first-wave follicle oocytes that pause their growth

**DOI:** 10.1101/2024.08.27.609995

**Authors:** Bikem Soygur, Eliza A. Gaylord, Mariko H. Foecke, Steven A. Cincotta, Tegan S. Horan, Anna Wood, Paula E. Cohen, Diana J. Laird

## Abstract

Ovulation results from the cyclical recruitment of non-renewing, quiescent oocytes for growth. Therefore, the primordial follicles that are established during development from an oocyte encapsulated by granulosa cells are thought to comprise the lifelong ovarian reserve ^1–4^. However, using oocyte lineage tracing in mice, we observed that a subset of oocytes recruited for growth in the first juvenile wave remain paused for many months before continuing growth, ovulation, fertilization and development into healthy offspring. This small subset of genetically-labeled fetal oocytes, labeled with Sycp3-CreERT2, is distinguished by earlier entry and slower dynamics of meiotic prophase I. While labeled oocytes were initially found in both primordial follicles and growing follicles of the first wave, they disappeared from primordial follicles by puberty. Unexpectedly, these first-wave labeled growing oocytes persisted throughout reproductive lifespan and contributed to offspring at a steady rate beyond 12 months of age, suggesting that follicles can pause mid-growth for extended periods then successfully resume. These results challenge the conclusion from lineage tracing of granulosa cells that first-wave follicles make a limited contribution to fertility^5^ and furthermore suggest that growth-paused oocytes comprise a second and previously unrecognized ovarian reserve.

## Main Text

Each of the ∼500 ovulations that occur in a female human’s lifetime is the culmination of a process that begins with millions of oocytes in the fetus and progresses through multiple developmental stages, including an arrest in meiosis and growth lasting up to 5 decades. From the large starting pool, the vast majority of fetal oocytes die during meiotic prophase and the subsequent encapsulation by pre-granulosa cells that forms primordial follicles, which are considered the lifelong ‘reserve’ of fertility. Of the small fraction of quiescent primordial follicle-residing oocytes recruited for growth, even fewer complete growth and maturation to reach ovulation^1–4^. Fertility and reproductive lifespan depend upon oocyte quantity and quality, but whether the process by which oocytes are selected for survival and growth to ovulation is stochastic or fitness-based, and the dynamics of growth stage progression remain unclear. As both quantity and quality of mature oocytes decrease with age, understanding the trajectory of oogenesis and the basis for selection would inform assisted reproductive treatments and fertility preservation.

As in humans, mouse oogenesis begins when primordial germ cells cease proliferation and enter meiotic prophase I (MPI); this transition occurs gradually from embryonic day (E)12.5 to E14.5^6^, first from the anterior center to periphery and then toward the posterior of the fetal ovary^7–10^. Oocytes proceed through MPI until the final diplotene stage, at which point meiosis arrests, and does not resume until the final stages of growth just before ovulation^11^. The primordial follicles have two developmental paths depending upon their initial location within the ovary. Primordial follicles that aggregate from pre-granulosa cells in the outer, cortical part of the ovary begin assembling before birth in mice^4,12^ and are completely formed by postnatal day 5 (P5)^13^. However, a subset of primordial follicles develop from pre-granulosa cells in the ovarian medulla and these begin growing beyond primordial stages immediately after formation. However, given this so-called ‘first wave’ of follicle growth precedes the production of gonadotropins required for maturation beyond the antral stage^8,14^, these prepubertal follicles are believed to die^15,16^. Lineage tracing of first wave follicles by labeling granulosa cells with *FoxL2-CreER* at E16.5^5^ showed they disappear by P90, suggesting temporally that the first wave of mature follicles recruited for growth could potentially contribute to the first litter conceived ∼P55 and born ∼P75 but with diminishing odds to later litters, although this is yet to be demonstrated. The majority of oocytes ovulated during reproductive life are believed to derive from the pool of quiescent primordial follicles in the ovarian cortex which are cyclically recruited for growth^16,17^. Histology as well as mutant phenotypes have shaped the hypothesis that follicle cohorts grow continuously through primary, secondary, and antral stages, after which gonadotrophin responsiveness promotes survival and growth of a subset in a process called follicle selection, while remaining antral follicles die^16,18^. However, we lack a detailed understanding of the developmental trajectory of first wave follicles and whether they contribute to female fertility.

Here, we used an alternative approach to track a cohort of the first wave follicles over time, by genetic labeling the early meiotic oocytes directly rather than the surrounding granulosa cells and thereby identified a subset of mouse fetal oocytes distinguished by earlier onset and slower progression through MPI. While these labeled oocytes initially demonstrated increased survival, and contributed to primordial follicles as well as the first wave of growing follicles in neonates, all of the labeled primordial oocytes disappeared by puberty. Labeled first wave oocytes unexpectedly persisted and gave rise to offspring at a steady rate throughout reproductive life. This lineage tracing and functional study of an oocyte subset from fetal development through advanced reproductive age demonstrates that, contrary to prior interpretations, the first wave of prepubertal growing follicles in mice can contribute to lifelong fertility by pausing their development for up to 11 months before resuming growth and ovulation.

## Results

### Meiotic prophase timing is heterogeneous and a unique subset can be lineage-traced

To characterize the spatio-temporal dynamics of MPI, we employed *in toto* immunostaining and optical clearing of intact mouse fetal ovaries. We previously identified the earliest meiotic entry in the anterior central region of the ovary at E12.5 (CD1 background), marked by the meiosis-specific synaptonemal complex protein 3 (SYCP3) in ∼5% of germ cells, which subsequently spread radially while sweeping posterior^10^. At E14.75, SYCP3 was present in all germ cells while SYCP1, which is recruited to the synaptonemal complex in second stage (zygotene) and is robustly expressed in the third stage (pachytene) of meiotic prophase I^19,20^, was largely undetectable in whole-mount stained ovaries. Rapid and simultaneous appearance of SYCP1 in all oocytes between E14.75 and E15.0 contrasted with the gradual onset of the first stage (leptotene) meiosis marker SYCP3 over two days. Other markers of meiotic progression turned on in a synchronous pattern including the synaptonemal complex central element protein-2 (SYCE2), which is required for completion of synapsis in the pachytene stage of MPI^21^ and HORMAD1, which is associated with unsynapsed chromosome axes^22^ (**Extended Data Fig. 1a, b**). At a population level, this discrete spatiotemporal patterning between the protracted onset of leptotene and synchronous progression to pachytene stage indicates that the time spent in MPI is not equal, with some oocytes dwelling longer in the early stages (**Fig. 1a**).

**Fig. 1.**
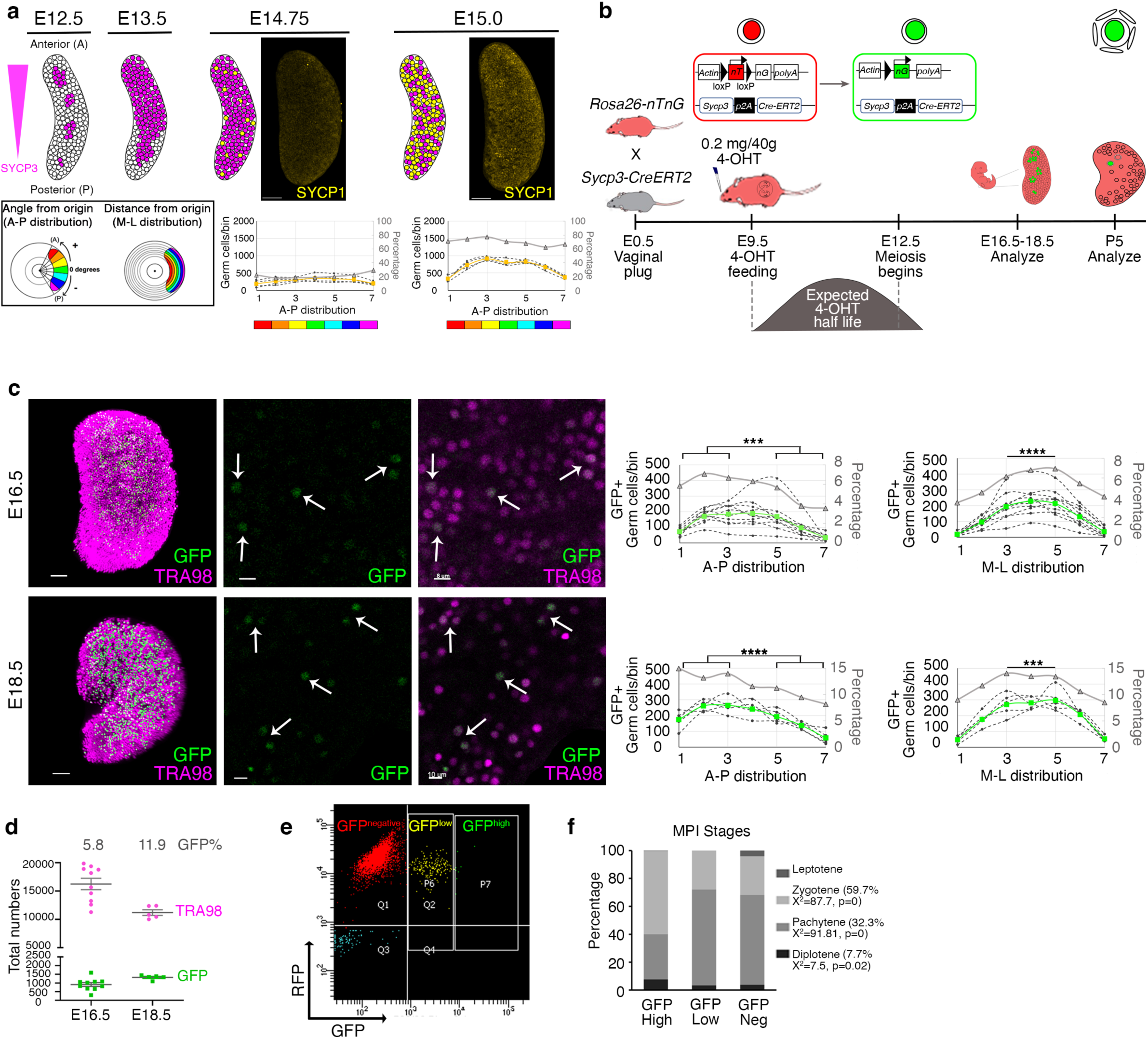
The earliest differentiating oocytes are distinct in their spatial distribution and meiotic progression. **a**, Schematic of earliest SYCP3 expressing cells (magenta) in the anterior-medial region of the E12.5 ovary, radiating outward by E14.75. For analysis, the ovary was divided into seven segments. Y-axes depict the number of germ cells in each bin, dashed lines represent individual ovaries, and color-coded bold lines indicate mean values for each antibody. Secondary y-axes (gray) show the percentage of cells positive for antibody of interest (i.e. SYCP1 in yellow) normalized to total germ cells in each bin. n=4 wild-type ovaries for each time point. Scale bars, 70 µm. **b,** Illustration of induced *in vivo* lineage tracing of the first meiotic entrants. Following 4-OHT feeding at E9.5, CreERT2 recombinase excises nT and initiates nG expression, causing constitutive expression of nuclear GFP in SYCP3-expressing cells. **c,** Whole-mount immunolabeling of TRA98 and GFP (white arrows) in *Sycp3^CreERT^*^2^*^/+^; Rosa26^nT-nG/+^* ovaries at E16.5 and E18.5. GFP labeled first meiotic entrants were predominantly located in the anterior (bins 1 to 3 on x axis of A-P distribution graph; \*\*\**P*=0.0005 for E16.5 and \*\*\*\**P*<0.0001 for E18.5) and middle (bins 3 to 5 on x axis of M-L distribution graph; ****P<0.0001 for E16.5 and ***P=0.0005 for E18.5) of E16.5 (n=10) and E18.5 (n=5) *Sycp3^CreERT^*^2^*^/+^; Rosa26^nT-nG/+^* ovaries. In graphs, distributions of GFP are shown in green. Scale bars are 50 µm for the whole ovary view, 8 µm for higher magnification at E16.5, and 10 µm for higher magnification at E18.5. **d**, Quantification of TRA98-positive total germ cells and GFP expressing first meiotic entrants at E16.5 (n=10) and E18.5 (n=7). **e,** FACS isolation of GFP^high^ GFP^low^ ,and GFP^negative^ cells from *Sycp3^CreERT^*^2^*^/+^; Rosa26^mT-mG/+^* ovaries at E17.5. **f,** MPI staging of chromosome spreads from nuclei of sorted cells in (e) following SYCP1/3 immunostaining (100 nuclei examined in each population, a total of n=900 cells were analyzed).

To determine the effect of MPI duration on oocyte fate, we performed *in vivo* lineage tracing of oocytes. We generated a drug-inducible *Sycp3-CreERT*2 knock-in mouse to mark germ cells committed to MPI. When crossed to *Rosa26^mTmG^*, which expresses tdTomato (RFP) in every cell, activation of the *Sycp3* promoter will cause excision of RFP and allow permanent and constitutive expression of GFP in those cells and their descendants. (**Extended Data Fig. 1c, d**). In a cross between *Sycp3^CreERT^*^2^*^/+^*males and *Rosa26^mTmG^* females, intraperitoneal Tamoxifen injected into pregnant dams at E11.5 (24 hours before the earliest meiotic initiation) led to GFP expression exclusively in oocytes throughout *Sycp3^CreERT^*^2^*^/+^; Rosa26^mTmG^* fetal ovaries by E14.5 with high efficiency (**Extended Data Fig. 1e, f**). To achieve precise quantification of labeled oocytes by *in toto* immunostaining of the ovary, we used the similar *Rosa26^nTnG^* reporter with nuclear GFP/RFP and TRA98 nuclear germ cell markers which facilitate cell segmentation during imaging^23,24^. We found that oral delivery of 4-hydroxytamoxifen (4-OHT), even at a lower dose, produced higher efficiency labeling of TRA98+ oocytes than intraperitoneal injection (**Extended Data Fig. 1g-i**) and confirmed that patterning and timing of MPI was normal (**Extended Data Fig. 1j**). However, administering low dose 4-OHT (0.2 mg/40 g) at E11.5 labeled over 50% of germ cells, owing to the prolonged effect of 4-OHT. (**Extended Data Fig. 1g-i**). To pulse label the earliest meiotic entrants observed at E12.5 while minimizing perdurance and later excision, we dosed with 4-OHT at E9.5 (**Fig. 1b**). At E16.5, GFP labeling was confined to 5.8% of TRA98+ germ cells in *Sycp3^CreERT^*^2^*^/+^; Rosa26^nTnG/+^* fetal ovaries (**Fig. 1c**); this frequency as well as the localization of labeled cells in the ovary (anterior/medial region) recapitulates our prior observation of endogenous SYCP3 at E12.5^10^. From an average 16,236 TRA98+ oocytes in E16.5 ovaries, 913 were GFP+, while at E18.5 1,457 were detected (**Fig. 1d and Supplementary Table 1**). Considering that meiotic oocytes do not proliferate, this modest increase in GFP could arise from residual 4-OHT or still accumulating GFP. To validate our labeling, we performed meiotic staging of oocytes based on their chromosomes. Using flow cytometry to separate GFP-labeled oocytes, we analyzed meiotic spreads from *Sycp3^CreERT^*^2^*^/+^; Rosa26^mT-mG/+^* ovaries at E17.5 following 4-OHT at E9.5 (**Fig. 1e, f and Extended Data Fig. 2a, b**). Whereas GFP^negative^ oocytes were primarily in pachytene stage as expected^25^, the GFP^high^ population showed a different distribution, with increased frequency in earlier zygotene (59.7%, Χ^2^ = 87.7, p=0.0) and later diplotene (7.7%, Χ^2^ = 7.5, p=0.02); a GFP^low^ population of oocytes matched the MPI stage profile of GFP^negative^ (**Fig. 1f**) and likely represents cells that were newly excised given their moderate levels of RFP (**Fig. 1e**). Together these results indicate that the MPI timing is heterogeneous among fetal oocytes, and that pulse-labeling with *Sycp3-CreERT*2 reveals distinct behavior of the earliest meiotic entrants.

### Enhanced survival of earlier meiotic entrants

From E16.5 to E18.5, we observed divergent behavior of the labeled oocytes. While the total number of TRA98+ oocytes declined by 25%, from a mean of 16,236 to 12,403 reflecting the normal process of fetal oocyte attrition (FOA^26–29)^, the number of GFP+ oocytes slightly increased (**Fig. 1d**). One possible explanation for increased survival during FOA could be a result of less meiotic recombination-associated DNA damage ^30–32;^ however the number of persistent RAD51+ unrepaired double-strand breaks at pachytene did not differ between GFP^high^, GFP^low^, and GFP^negative^ E17.5 meiotic spreads (**Fig. 2a**). Longer MPI duration for earlier entrants could affect the formation of crossovers between meiotic chromosomes, and could also explain survival enhancement as failures in synapsis lead to oocyte elimination^33^. However, the number of MLH1+ foci per pachytene nucleus was also unchanged (**Fig. 2b and Extended Data Fig. 2c**). Direct examination of chromosome synapsis via SYCP3 and SYCP1 colocalization revealed that the fraction of nuclei with fragmented synaptonemal complexes was increased in GFP^high^ and GFP^low^ compared to GFP^negative^ and there was a trend toward elevated partial asynapsis (**Fig. 2c and Extended Data Fig. 2d**). These analyses suggest that meiotic fidelity may be compromised in the earliest meiotic entrants, but normal levels of double-strand break repair do not explain the enrichment of GFP+ oocytes during FOA.

**Fig. 2.**
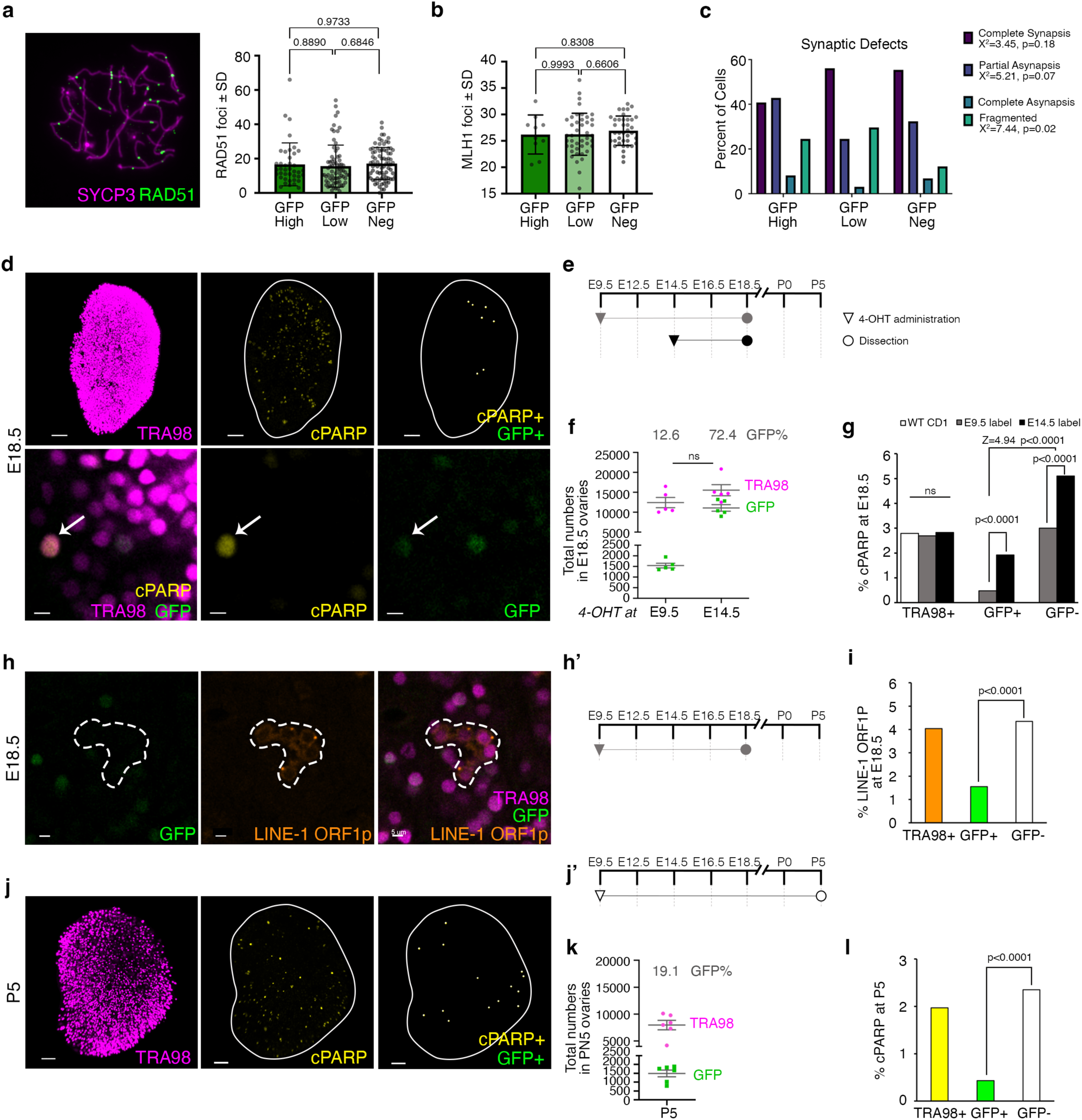
Sycp3-CreERT2 labeled early meiotic entrants are more resistant to FOA. Quantification at E17.5 of (**a**) persistent double-strand breaks with RAD51 (n=40 nuclei for GFP^high^, n=74 for GFP^low^, and n=81 for GFP^negative^), (**b**) class I crossovers with MLH1 (n=10 nuclei for GFP^high^, n=43 for GFP^low^, and n=38 for GFP^negative^), and (**c**) synaptic defects with SYCP3/1 (n=49 nuclei for GFP^high^, n=98 for GFP^low^, and n=81 for GFP^negative^). **d,** Whole-mount immunolabeling of cells expressing TRA98, cPARP, and GFP in E18.5 *Sycp3^CreERT^*^2^*^/+^; Rosa26^nT-nG/+^* ovaries following E9.5 4-OHT administration. White lines delineate ovary borders and white arrows indicate a cPARP+ GFP+ TRA98+ oocyte. Scale bars are 50 µm for whole ovary view (top) and 10 µm for higher magnification (bottom). **e,** Timeline for 4-OHT administration and evaluation. **f,** Quantification of TRA98+ (magenta) and GFP+ cells in E18.5 *Sycp3^CreERT^*^2^*^/+^; Rosa26^nT-nG/+^* ovaries following 4-OHT at E9.5 (labeling first meiotic entrants, n=5 ovaries) and E14.5 (random oocyte labeling, n=5 ovaries). The number of TRA98+ oocytes in each labeling condition was similar by T-test. **g,** Frequency of cPARP colocalization with TRA98+ oocytes, GFP+ and GFP-negative oocyte subsets at E18.5 following 4-OHT dosing at E9.5 or E14.5. Frequencies by labeling time were compared by Chi-square test whereas the p-value for the interaction term of 4.94 computed between 4-OHT timing, cPARP and GFP is shown above with Z-statistic. **h,** Whole-mount immunostaining and (**i**) quantification of GFP, LINE-1 ORF1p and TRA98 in E18.5 *Sycp3^CreERT^*^2^*^/+^; Rosa26^nT-nG/+^* ovaries following (**h’**) 4-OHT administration at E9.5. Scale bars, 5 µm. LINE-1 ORF1p expression was predominantly observed in GFP-TRA98+ oocytes (white dashed lines) compared to GFP+ TRA98+ oocytes. **j,** Whole-mount immunostaining of TRA98, cPARP, and GFP in P5 *Sycp3^CreERT^*^2^*^/+^; Rosa26^nT-nG/+^* ovaries following (**j’**) 4-OHT at E9.5. Scale bars, 50 µm. **k** and **l,** Quantification of TRA98 and GFP+ cells revealed lower frequency of cPARP positivity in GFP+ compared to GFP-negative oocytes.

We examined apoptosis directly in different subpopulations of fetal oocytes by immunostaining intact ovaries with cleaved PARP (cPARP; **Fig. 2d and Extended Data Fig. 2e**). At E18.5, we observed nuclear cPARP in ∼3% of TRA98+ fetal oocytes in untreated CD1 mice as well as in mixed background *Sycp3^CreERT^*^2^*^/+^; Rosa26^nT-nG/+^* mice with gestational 4-OHT dosing (**Fig. 2e-g**) and in both, apoptosis in the earliest meiotic entrants occurred without spatial bias (**Extended Data Fig. 2f**). After 4-OHT at E9.5, GFP+ oocytes comprised 12.6% of all TRA98+ oocytes at E18.5, but only 0.46% co-labeled with cPARP in the ovaries analyzed for apoptosis(**Fig. 2d-g and Extended Data Fig. 2e)**. To test the possibility that GFP expression affects the threshold of apoptosis in oocytes, we shifted 4-OHT administration to E14.5 (**Fig. 2e**), when oocytes ubiquitously express SYCP3^10^. After this late and random labeling, GFP+ oocytes comprised 72.4 % of all TRA98+ oocytes at E18.5 (**Fig. 2f**), but 1.9% co-labeled with cPARP, compared to the 2.8% overall rate of apoptosis in all cPARP+ TRA98+ oocytes (**Fig. 2g**). By chi-square test, the frequency of cPARP+ oocytes in the GFP+ versus GFP-negative compartments differed between early and late 4-OHT dose (P<0.001) (**Fig. 2g**). However, as a threeway measure of the association between cPARP and GFP, we computed the odds ratio in each 4-OHT labeling experiment as 2.34 and a Z-statistic for this interaction term of 4.94 (P<0.0001), indicating a strong difference in the relationship between cPARP and GFP in early versus late labeling experiments (**Fig. 2g**). These analyses demonstrate that in the *Rosa26^nT-nG^* mouse, GFP expressing fetal oocytes are protected against apoptosis compared to RFP+ oocytes, while the overall rate of apoptosis is unchanged. Beyond this important caveat about fluorescent reporters in fetal oocytes, our studies support a reduced frequency of apoptosis at E18.5 in *Sycp3-CreERT2*-labeled early meiotic entrants.

Since the activity of the LINE-1 (L1)–transposable element has been functionally linked to FOA in mice^34,35^, we tested the possibility that GFP+ early meiotic entrants express L1 ORF1p at lower levels. At E18.5, 4.0% of TRA98+ oocytes expressed L1 ORF1p, compared to only 1.9% of GFP+ oocytes (p<0.0001; **Fig. 2h-i**). This result further substantiates mitigated FOA in the earliest meiotic entrants.

As a substantial proportion of oocytes are eliminated during cyst breakdown and formation of primordial follicles^4^, we examined *Sycp3^CreERT^*^2^*^/+^; Rosa26^nT-nG/+^* ovaries at P5, when follicle formation is largely complete (**Fig. 2j and j’**). While the average number of TRA98+ oocytes fell from 12,403 at E18.5 to 7,959 at P5 (**Supplementary Table 1**), the number of early labeled GFP+ oocytes remained constant (a mean of 1,485 compared to 1,457 at E18.5; **Fig. 1d**, **2k**). An increased number of TRA98+ oocytes underwent apoptosis in the cortex of the P5 ovary, with a notable absence of cPARP in the medullary follicles (**Extended Data Fig. 2g, h**). While the frequency of cPARP in TRA98+ oocytes at P5 was 2.0%, there was a 4-fold cPARP reduction in the GFP+ subpopulation (**Fig. 2l).** This decreased apoptosis, together with the successive enrichment of GFP+ oocytes from 5.8% at E16.5 to 19.1% at P5 (**Fig. 1d and 2k**), corroborates a survival advantage of the earliest meiotic entrants during FOA and follicle formation.

### Elimination of labeled primordial, but not growing, follicles by puberty

The first wave of follicle growth immediately follows follicle formation^5,8,36^, but the relationship between meiotic entry, follicle formation and first-wave growth remains unclear. In *Sycp3^CreERT^*^2^*^/+^; Rosa26^nT-nG/+^* ovaries at P5, we identified that approximately 3% of total oocytes are actively growing using two criteria: decreased expression of TRA98 ^37^ together with the presence of AMH secreted by surrounding granulosa cells^38^(**Fig. 3a and Supplementary Movie 1**). *Sycp3-CreERT*2 lineage-traced GFP+ oocytes displayed no spatial bias following the compartmentalization of the cortex and medulla (**Extended Data Fig, 3a**). The frequency of GFP+ oocytes in growing follicles at P5 was 25.3%, similar to 19.1% GFP+ oocytes in primordial/primary follicles in the surrounding cortex (**Fig. 3b, c**). Contrary to the Production Line Model that posits that oocyte developmental order is determined by follicle formation order^15,39,40^, this observed equal partitioning of labeled oocytes between the first wave of growing medullary follicles and the non-growing follicles suggests that initial follicle compartmentalization is not related to timing of meiotic entry.

**Fig. 3.**
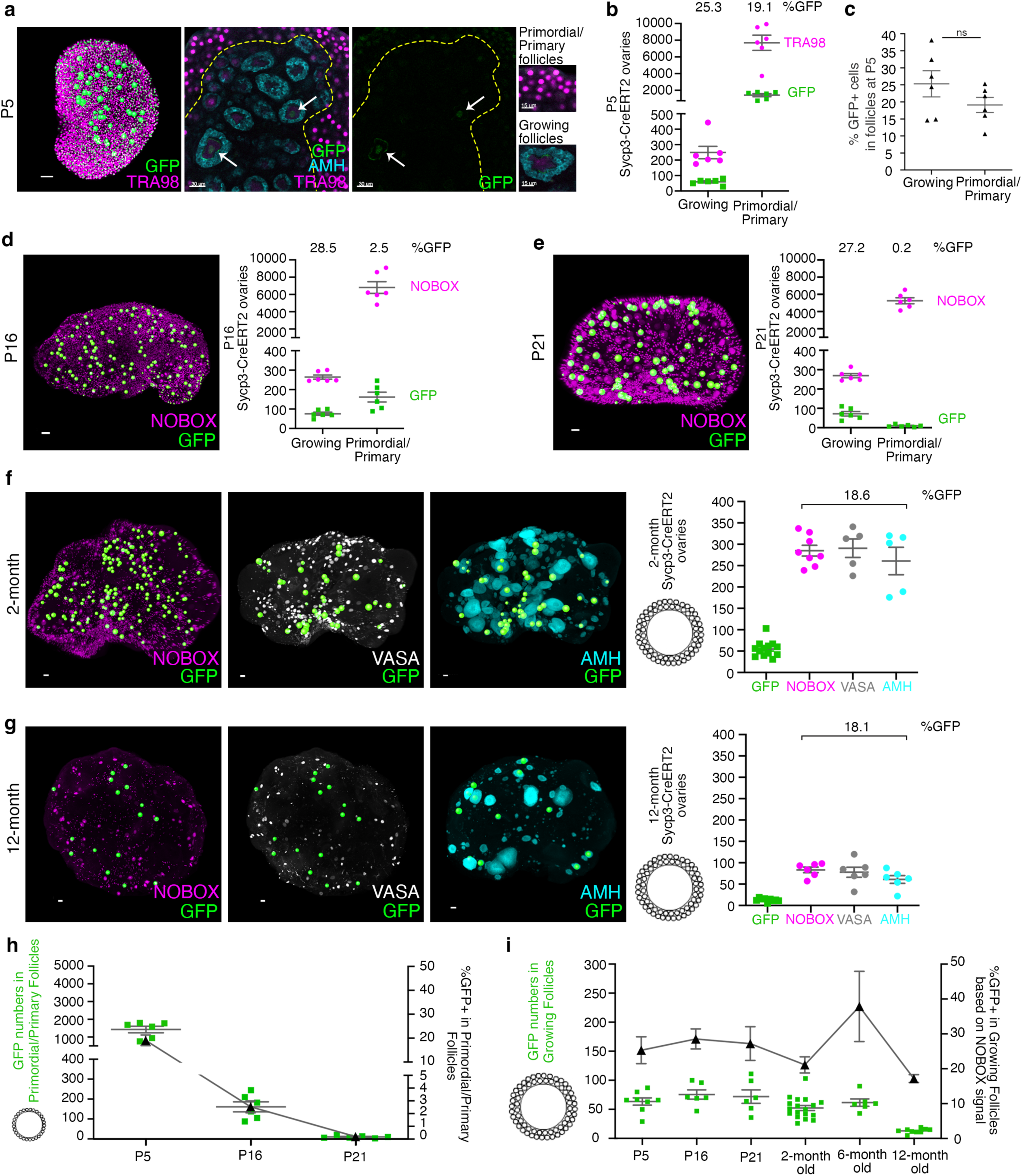
GFP expression from Sycp3-CreERT2 disappears from primordial/primary follicle oocytes by puberty but persists in growing follicle oocytes for up to 12 months. **a**, Whole-mount immunostaining and (**b**) quantification of growing follicles using AMH (granulosa cell marker) expression and low TRA98 expression in *Sycp3^CreERT^*^2^*^/+^; Rosa26^nT-nG/+^* ovaries (n=6) revealed a comparable percentage of GFP+ oocytes (**c**) in both growing and primordial/primary follicles at P5. Ovarian medulla is outlined by the yellow dashed line, with white arrows indicating AMH+ growing follicles containing GFP+ oocytes. The scale bars are 50 µm for the whole ovary view (left), 30 µm for the higher magnification (middle), and 15 µm for the primordial/primary and growing follicles (right). **d,** At P16 and (**e**) P21, whole-mount immunostaining against oocyte marker NOBOX and GFP showed a dramatic reduction in primordial/primary follicles containing GFP+ oocytes in the cortex (n=6 for each timepoint). Scale bars, 50 µm. Contribution of GFP+ oocytes to the dynamic pool of growing follicles (**f**) in 2-month (n=13) and (**g**) 12-month-old (n=9) *Sycp3^CreERT^*^2^*^/+^; Rosa26^nT-nG/+^* ovaries, assessed using NOBOX and VASA (oocyte markers) and AMH (granulosa cell marker). Scale bars, 50 µm. Each data point in b, d, e, f, and g corresponds to an individual ovary and the percentage of GFP+ oocytes are displayed at the top of the graphs. Quantifications of GFP+ oocytes (**h**) in primordial/primary (P5 to P21) and (**i**) growing (P5 to 12-month) follicles highlighted the disappearance of GFP+ primordial/primary follicles, while GFP+ oocytes in growing follicles were observed until 12-months. The primary y-axes display the total number of GFP+ oocytes, while the secondary y-axes represent the percentage of GFP+ oocytes within the total oocyte population. The x-axes indicate various time points.

During postnatal development, we observed a precipitous decline in the total number of GFP+ oocytes in *Sycp3^CreERT^*^2^*^/+^; Rosa26^nT-nG/+^* ovaries: from a mean of 1,485 at P5, to 237 at P16 to 82 at P21 (**Supplementary Table 1**). In contrast to the initial equivalence of labeled oocytes in the growing and non-growing (primordial/primary) follicle compartments at P5, GFP+ growing oocytes remained constant in number and frequency from P5 to P21 while GFP+ primordial/primary oocytes decreased nearly 100-fold from 19.1% at P5 to 0.2% at P21; this loss of GFP+ primordial follicle oocytes far exceeds the overall 32% reduction in the total primordial follicle pool during this period (**Fig. 3b-e, Extended Data Fig. 3b, c**, and **Supplementary Movies 2 and 3**). Beyond P21, when an average of 9.8 GFP+ primordial/primary oocytes was identified throughout the entire ovary, the number became so rare that we ceased to quantify. Although juvenile oocyte attrition is not well understood, the accelerated rate of elimination of GFP+ oocytes from primordial/primary follicles during the prepubertal period contrasts with the enhanced survival of GFP+ oocytes during FOA. These temporally disparate fates of labeled cells argue that the process of oocyte selection operates by different criteria during the fetal and juvenile periods.

While primordial/primary GFP+ oocytes disappeared by puberty, growing GFP+ oocytes remained surprisingly constant in number and frequency from P5 to P21 (**Fig. 3d-e**). To identify the transcriptional signature of GFP+ oocytes, we performed bulk RNA sequencing. Following superovulation, we collected MII oocytes at P21. Bulk RNA-seq analysis revealed that the transcriptomes of superovulated GFP+ and GFP-/RFP+ oocytes were remarkably similar, with only 12 differentially expressed genes (**Extended Data Fig. 4a-g**). Together these results suggest that the GFP+ oocytes that reach maturity are transcriptionally equivalent, and that the consequences of meiotic timing are likely limited to selection of fetal oocytes and prepubertal primordial follicles.

### Labeled oocytes in growing follicles persist through adulthood

Since previous work showed that pulse-labeled granulosa cells in neonatal growing follicles disappear by 3 months^5,36,41^, we might expect the dynamics of labeled oocytes in the same first-wave of growth to be similar. However, young adult mice at 2 months of age still retained an average of 55.2 GFP+ oocytes per ovary (**Fig. 3f and Supplementary Movie 4**). Co-immunostaining with four markers verified that all GFP+ oocytes were in early growth: AMH (secreted by granulosa cells from small growing follicles^42,43^), NOBOX (nuclear and increasingly in the cytoplasm of primary and secondary oocytes^44^), VASA (in all stages of oocytes,^4^ distinguished by volume) (**Fig. 3f and Extended Data** Fig. 3d), and GDF9 (in oocytes from secondary stage^45^; **Extended Data Fig. 3e**). At 6 months of age, we detected an average of 61.7 GFP+ growing oocytes per ovary (**Extended Data Fig. 3f**), declining to 13.0 at 12 months (**Fig. 3g and Extended Data** Fig. 3g). Since GFP+ primordial/primary oocytes declined rapidly between P5 and P21 (**Fig. 3h**) the GFP+ growing oocytes observed at steady numbers until 6 and 12 months of age (**Fig. 3i**) could not have arisen by cyclic recruitment from the primordial follicle pool. Rather, the persistence of a small number of labeled growing oocytes for 11 months after the disappearance of labeled primordial/primary oocytes suggests that, once committed to growth, oocytes have the potential to arrest and survive without undergoing atresia.

### GFP+ offspring are consistently produced throughout adulthood

We next evaluated the capacity of the GFP+ growing oocytes in adult ovaries to generate offspring. Upon reaching sexual maturity, 14 *Sycp3^CreERT^*^2^*^/+^*; *Rosa26^nT-nG/+^*females that received 4-OHT *in utero* at E9.5 were housed with wild-type males. Although all oocytes were phenotypically either GFP+ or RFP+, only half of progeny should inherit the *Rosa26^nT-nG^*allele and express either reporter ubiquitously (**Fig. 4a**). Among 1,134 pups screened at P1-3, 51.4% of the progeny carried one fluorescent protein, indicating that neither GFP nor RFP affects the competence of oocytes or development of embryos (**Fig. 4b**). As the breeding females aged, the fraction of GFP+ pups remained consistent until the tenth litter (**Fig. 4c**) even though overall fecundity declined after 9 months (**Fig. 4d-f**). Over 8 months, an average of 95.3 pups were beget by each female (38.5 RFP+ and 10.1 GFP+) with equal frequency across time (**Fig. 4g**). The observed incidence of GFP in 17-37% of growing oocytes within the ovary at 2, 6 and 12 months (**Fig. 3f, g and Extended Data** Fig. 3e, f) predicts that half, or 9-19% of offspring would be GFP+ if maturation and fertilization were stochastic. In agreement, the constant 10.6% rate of GFP+ pups obtained per female demonstrates that follicles are competent after prolonged growth arrest, and indicates that the first meiotic entrants in the fetal ovary contribute durably to offspring throughout reproductive life.

**Fig. 4.**
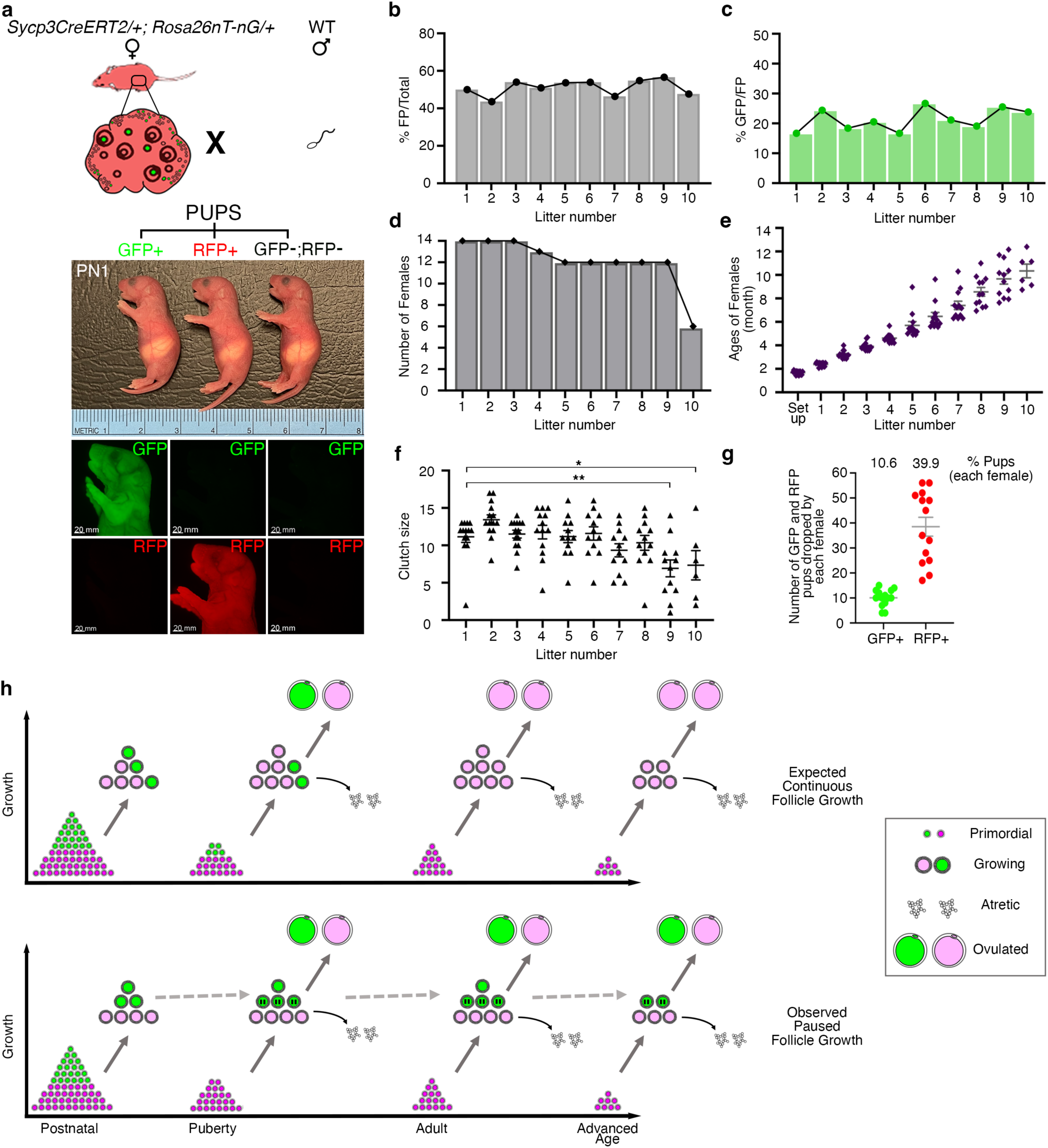
Sycp3-CreERT2 oocytes labeled at the onset of meiosis contribute to offspring at a constant frequency throughout reproductive lifespan. **a**, *Sycp3^CreERT^*^2^*^/+^; Rosa26^nT-nG/+^* females exposed to 4-OHT at E9.5 (n=14) were crossed with wild-type males and pups were screened to detect embryos derived from GFP+ oocytes (left), RFP+ oocytes (middle) or GFP-;RFP-unlabeled oocytes (right). **b,** Percentage of all pups expressing fluorescent protein (FP). **c,** Percentage of GFP+ pups among FP-expressing pups remained similar across litters despite increasing maternal age. **d,** Half of females ceased reproducing after the ninth litter. **e,** Ages of the females and (**f**) the number of pups produced per female in each litter. The number of pups per litter decreased in the 9th (**P=0.0038) and 10th (*P=0.0381) litters compared to the first litter. **g,** Cumulative number of GFP+ and RFP+ pups produced by each female (circles, with mean indicated as gray line) upon reaching 12 months of age. **h,** Model of paused follicle growth in mouse ovaries; (top) the paradigm of continuous follicle growth in which growing follicles are cyclically recruited from primordial follicles and either undergo atresia or reach ovulation stage which predicts that GFP-labeled first-wave growing oocytes should be rapidly depleted, with ovulation in adulthood and later reproductive life resulting from continuous recruitment from unlabeled primordial follicles. However, detection of GFP+ pups up to 12 months of age, alongside persistence of GFP labeled growing oocytes/the loss at puberty of GFP labeled primordials, leads to the model in which growing follicles in the medulla derived from the first meiotic entrants can pause, and later resume growth and maturation to contribute to offspring until 12-months (bottom).

## Discussion

This study reveals heterogeneity in the developmental trajectories of oocytes, during MPI and follicle growth. Our system of pulsed genetic labeling of fetal oocytes together with *in toto* imaging of the entire optically cleared ovary enabled precise quantification of labeled postmitotic oocytes throughout postnatal development, puberty, aging, and assessment of their developmental competence. These breeding studies functionally demonstrate that the first wave of follicle growth contributes to lifelong fertility.

The persistence of labeled oocytes from first wave follicles— in contrast to the disappearance of labeled granulosa cells^5,36,41^— indicates that the somatic compartment of preantral follicles is more dynamic than previously appreciated. These divergent results raise the possibility that granulosa cells in first-wave follicles are replaced over time by other somatic support cells. Given that only growing GFP+ oocytes remain beyond puberty, the sustained derivation of GFP+ offspring is incompatible with the paradigm that primordial follicles are the sole source of mature, ovulated oocytes throughout life. Considering that it typically takes 6-12 days for a secondary follicle to reach the antral stage^46^, the presence of GFP+ growing oocytes in the medulla more than 11 months after their disappearance from cortical primordial follicles by P21 suggests that these GFP+ follicles remained in the ovary for over 35 cohorts of follicle growth before giving rise to GFP+ offspring. Based on these results, we propose that oogenesis in mice also occurs through an alternative trajectory in which prepubertal growing follicles pause for variably extended periods before resuming growth, maturation, and ovulation (**Fig. 4h**). Stable isotype labeling of postnatal mouse ovaries revealed the persistence of long-lived proteins (and specifically ZP3) in primary and secondary follicles until 6 months of age^47^. Given that ZP3 is uniquely expressed in oocytes during their growth and the labeling pulse was administered during postnatal development, these results support our model of growing follicle pause.

Our observations in ovaries from 2 to 12 months suggest that the majority of putative paused follicles are at the secondary stage (**Extended Data Fig. 3d, g**). Given that the pausing was revealed by pulse-labeling a fetal oocyte subpopulation that only gives rise to the first wave of growing follicles, we cannot draw any conclusions on the potential for later waves of recruited follicles to pause.

Our data further suggest that there is no penalty for pausing growth. Congruence between the observed frequency of GFP+ presumptive paused follicles in adult ovaries and GFP+ offspring born at a constant rate throughout life demonstrates that labeled and unlabeled oocytes are equally likely to complete growth, ovulation, fertilization and embryonic development. However, in spite of the observed equivalence of births from paused GFP+ follicles and from unlabeled oocytes presumably directly recruited from the primordial follicle pool, there may be important consequences for the two trajectories. Given the differences in metabolism and repair in primordial versus preantral oocytes^48,49^, the rate of aneuploidy and competence may diverge at advanced maternal age for oocytes that have undergone pausing compared to those directly recruited from primordial follicles.

The finding that follicle growth is not necessarily continuous but can pause could have significant relevance for fertility. If pausing of follicle growth occurs in humans, this would constitute a second ovarian reserve in addition to the known primordial follicle reserve. The ability to manipulate follicle growth pausing would potentially expand options for fertility treatment and the extension of reproductive lifespan.

## Methods

### Mice

Engineering of the Sycp3-CreERT2 construct was carried out by Bacpac Genomics. To generate a knock-in mouse line carrying an iCreERT2 driven by a functionally active mSycp3, the stop codon of mSycp3 was converted to P2A sequence. The P2A self-cleaving peptide ^50^ requires translation of SYCP3 for cleavage and nuclear localization of CreERT2 and ensures that the regulation of *Sycp3* and function of SYCP3 are maintained. The CRISPR/Cas9 system was utilized for aiding recombination and precise targeting. sgRNAs were designed using the CRISPR guide–design tool developed by the Zhang laboratory at MIT (crispr.mit.edu) for the Cas9 enzyme from *S. pyogenes* (PAM: NGG). This tool screens candidate sgRNAs for selectivity; it ranks conceivable off-target locations based on the number and location of mismatches relative to the sgRNA. The activity levels of this sgRNA were also checked for predicted activity level with a set of tools available on Crispr.org with abbreviated display on the UCSC Mouse Genome Browser.

The construct (carrying iCre) was modified by opening it with XhoI, which cuts very close to the end of iCre, and then inserting the PCR amplified ERT2 module. The PCR products included (in one PCR primer) the proper end of iCre leading to the repair of iCre (minus the stop-codon) and connection with ERT2. The cloned ERT2 module was sequenced in both orientations to ensure that the sequence matched the design.

Superovulated female FVB/N mice (4-weeks-old) were mated to FVB/N stud males, and fertilized zygotes were collected from oviducts. Cas9, sgRNA, and plasmid vectors were mixed and injected into the pronucleus of fertilized zygotes. After the injection procedure, zygotes were implanted into oviducts of pseudopregnant CD1 female mice. Mice positive for targeting were identified by genotyping. The CRISPR/Cas9 mediated targeting of iCreERT2 in FVB zygotes yielded 5 of 25 founder pups (20%) positive for targeting as indicated by genotyping. We outcrossed *Sycp3^CreERT^*^2^ positive (*Sycp3^CreERT^*^2^*^/+^*) males with CD1 females to remove possible off-target editing events.

The specificity and efficiency of the *Sycp3-CreERT2* line were tested by crossing *Rosa26^mT/mG^* ^51^ females with *Sycp3^CreERT^*^2^*^/+^* males. *Rosa26^mT/mG^* (MGI:J:124702) mice were outcrossed with CD1, and maintained on a mixed genetic background. Tamoxifen was injected intraperitoneally into pregnant *Rosa26^mT/mG^* dams at E11.5 at a concentration of 4 mg (Sigma-Aldrich, concentration of working solution; 40 mg/ml in sunflower seed oil) per 20 g of body weight for embryonic ovary collection at E14.5. To enhance the segmentation of labeled oocytes during analysis we used the *Rosa26^nT/nG^* line, which has nuclear localization of constitutive tdTomato and floxed GFP. *Rosa26^nT/nG^* females were crossed with *Sycp3^CreERT^*^2^*^/+^*males and 0.2 mg of 4-OHT (Sigma-Aldrich, concentration of working solution; 5 mg/ml in peanut oil) per 40 g of body weight was pipette fed to pregnant *Rosa26^nT/nG^* dams at E11.5. *Rosa26^nT/nG^* mice (MGI:J:199711) were purchased from the Jackson Laboratory, outcrossed with CD1, and maintained on a mixed genetic background. Embryonic ovaries were collected at E18.5, fluorescent signal was checked under a fluorescent dissection microscope (Olympus MVX10), and then processed for whole-mount immunofluorescence staining, resulting in the labeling of more than 50% of oocytes. To label the precise population of the first meiotic entrants in embryonic ovaries, *Rosa26^nT/nG^*females were crossed with *Sycp3^CreERT^*^2^*^/+^*males and 4-OHT (Sigma-Aldrich, concentration of working solution; 5 mg/ml in peanut oil) at a dose of 0.2 mg per 40 g body weight was pipette fed to pregnant dams at E9.5. Embryonic ovaries were collected at different stages of pregnancy (E13.5, E16.5, and E18.5).

For the preparation of meiotic spreads, *Rosa26^mT/mG^* females were crossed with *Sycp3^CreERT^*^2^*^/+^* males and 0.2 mg of 4-OHT per 40 g of body weight was pipette fed to pregnant dams at E9.5 for embryonic ovary collection at E17.5. To overcome dystocia in pregnant females after 4-OHT administration, pregnant dams were pipette fed 0.1 mg of progesterone (Sigma, P-3972) per 40 g of body weight (dissolved in peanut oil) at E16.5. Postnatal and adult ovaries were collected at P5, P16, P21, 2-month, 6-month, and 12-month and processed further for whole-mount immunofluorescence staining. All ovaries assessed in this study were from co-housed sexually-inactive females.

The *Sycp3-CreERT2* mouse line was generated at the Children’s Hospital of Oakland and Gladstone Institutes mouse core. All other mouse work was performed under the University of California, San Francisco (UCSF), Institutional Animal Care and Use Committee guidelines in an approved facility of the Association for Assessment and Accreditation of Laboratory Animal Care International. To determine the timing of pregnancies, female mice were set up with individual males and checked every morning for the appearance of the vaginal mating plug (vaginal plug). The day of the mating plug was identified as E0.5, and female mice were euthanized at different stages of pregnancy. CD1 female mice (purchased from Charles River) were mated with male mice to map the later stages of meiotic prophase I (MPI) in wild-type ovaries.

### Genotyping

Total genomic DNA was extracted from ear punches or tail tips by boiling in alkaline lysis reagent (25 mM NaOH ; 0.2 mM EDTA) for 45 minutes at 95°C, cooling to 4°C, and then neutralizing with an equal volume of neutralization buffer (40 mM Tris-HCl). Primer sets used in the study are listed (**Supplementary Table 2**). For *Sycp3-CreERT2* genotyping, junction PCR was performed at the 5’ and 3’ ends, with each PCR spanning the “junction” between the inserted cassette and endogenous genomic sequence outside of the homology arms used for recombination. *Sycp3-CreERT2* Upstream (US) PCR was performed at 94°C for 2 min; followed by 10 cycles of 94°C for 10 sec, 55°C for 30 sec, 68°C for 90 sec; 20 cycles of 94°C for 10 sec, 55°C for 30 sec, 68°C for 90 sec (+ 20 sec per cycle); and 72°C for 5 min. *Sycp3-CreERT2* Downstream PCR was performed at 98°C for 30 sec; followed by 35 cycles of 98°C for 10 sec, 56°C for 15 sec, 72°C for 2 min; and 72°C for 2 min. *Rosa26-nTnG* and *Rosa26-mTmG* reactions were carried out as follows: 94°C for 2 min; followed by 10 cycles of 94°C for 20 sec, 65°C for 15 sec (-0.5°C decrease/per cycle), 68°C for 30 sec; 28 cycles of 94°C for 15 sec, 60°C for 15 sec, 72°C for 10 sec; and 72°C for 2 min. PCR products of *Sycp3-CreERT2* upstream and downstream reactions were electrophoretically separated in a 1% agarose gel and *Rosa26-nTnG* and *Rosa26-mTmG* reactions were separated in a 2% agarose gel. Gels were prepared with a tris/borate/EDTA buffer. Genotypes of adult mice and embryos were identified according to size of PCR products (**Supplementary Table 2**).

### Whole-mount immunofluorescence staining

Mouse fetal, postnatal, adult, and aged ovaries for whole-mount staining were dissected in 0.4% bovine serum albumin (BSA) in 1X phosphate buffered saline (PBS) and transferred into 2mL Eppendorf tubes. Subsequent steps were carried out while rocking as previously described ^23^.

Fetal (E13.5, E16.5, E18.5) and P5 ovaries were fixed with 4% paraformaldehyde (PFA) in PBS at +4°C for 2 hours then washed three times with 0.2% BSA in PBS for 10 min each. Ovaries were blocked with 2% BSA and 0.1% Triton X-100 in PBS for 3 hours at room temperature. Primary antibodies (**Supplementary Table 2**) were diluted in 0.2% BSA and 0.1% Triton X-100 in PBS, and ovaries were incubated in primary antibodies at +4°C for 5 nights. Samples were washed four times with 0.1% Triton X-100 in PBS for 15 min each at room temperature and incubated with Alexa Fluor–conjugated secondary antibodies in 0.2% BSA and 0.1% Triton X-100 in PBS at +4°C for 5 nights. Ovaries were washed three times with 0.2% BSA and 0.1% Triton X-100 in PBS for 30 min each, dehydrated with a methanol:PBS series (25 to 50 to 75 to 100%) for 10 min each (only 100% twice) at room temperature and incubated with 3% H2O2 in methanol overnight at +4°C. The following day, ovaries were incubated in 100% methanol for 30 min twice, transferred to a sample holder consisting of 10-mm-long glass cylinders (ACE Glass 3865-10) mounted onto coverslips (Fisherfinest Premium Cover Glass 12–548-5P) with silicone glue, then incubated in benzyl alcohol:benzyl benzoate (1:2) (BABB) at +4°C overnight. For imaging, mouse ovaries were oriented such that the anatomically anterior and posterior sections of the ovary were at the top and bottom of the field of view, respectively. Samples were imaged using a white-light Leica TCS SP8 inverted confocal microscope with a Fluotar VISIR 25X/0.95 water objective,1024 x 1024 pixel resolution, and stacks were acquired every 2 µm.

Postnatal (P16, P21) and adult (2-month, 6-month) ovaries were fixed with 4% paraformaldehyde (PFA) in PBS at +4°C for 3 hours while aged (12-month) ovaries were fixed for 4 hours. Postnatal, adult, and aged ovaries were washed four times with PBS for 20 min each. Ovaries were blocked with 0.2% Gelatin and 2% Triton X-100 in PBS overnight at +4°C. Primary antibodies (**Supplementary Table 2**) were diluted in 0.2% Gelatin, 2% Triton X-100, and 0.1% Saponin in PBS, and postnatal and adult ovaries were incubated in primary antibodies at +37°C for 1 week while aged ovaries were incubated in primary antibodies for 2 weeks. Samples were washed four times with 0.2% Gelatin and 2% Triton X-100 in PBS for 15 min each at room temperature. Samples were incubated with Alexa Fluor–conjugated secondary antibodies in 0.2% Gelatin, 2% Triton X-100, and 0.1% Saponin in PBS at +37°C for 3 nights followed by 5 days at +4°C. Ovaries were washed six times with 0.2% BSA and 0.1% Triton X-100 in PBS for 30 min each. Ovaries were incubated in an ascending tetrahydrofuran (THF): dH2O series (50% overnight at room temperature, followed by 80% then 100% THF for 1.5 hours each). To remove lipids, ovaries were incubated in dichloromethane (DCM) for 30 min at room temperature. For the final clearing step, ovaries were incubated in dibenzyl ether (DBE) at room temperature until they became transparent (in postanal samples, overnight clearing was sufficient, whereas in adult and aged ovaries at least 2 nights was required). Samples were imaged using a white-light Leica TCS SP8 inverted confocal microscope with a HC PL APO CS 10X/0.40 dry objective, 1024 x 1024 pixel resolution, and stacks were acquired every 2µm.

### Image analysis for whole-mount immunofluorescence stained ovaries

Image analysis was performed using Imaris software v8.3.1 (Bitplane) as previously described ^10,23^ with modifications. Samples were imported to Surpass mode, anterior and posterior parts of the ovary were defined according to orientation of attached mesonephros for embryonic samples and oviduct/uterus for postnatal, adult, and aged ovaries. A surface was created manually on the ovary and surrounding tissues were removed by masking the channels that corresponded to different antibody staining. Subsequent analysis was carried out using only the masked region of the ovary.

Fluorescently labeled germ cells were selected by using the Spot detection module. We utilized an object size criterion (XY diameter of 4 µm) to identify total germ cells based on TRA98 signal at E13.5, E16.5, E18.5. For P5 ovaries, oocytes within primordial follicles were identified as having an XY diameter of 4 µm while oocytes within growing follicles (in the medulla region) of P5 ovaries were manually chosen, considering their diminished TRA98 expression and larger XY diameter. We note that this selection method also captures oocytes in transitional and primary follicles. To count SYCP3, SYCE2, and HORMAD1 positive germ cells at E13.5, we applied object sizes of 3µm, 4.5 µm, and 4 µm, respectively. GFP-labeled cells at E16.5, E18.5, and P5 were identified through a semi-automated analysis: an object size filter of 4 µm was applied, followed by a manual analysis of the entire ovary to ensure accurate counting of GFP+ objects. To conduct apoptosis analysis, c-PARP+ cells were quantified using an object size filter of 4 µm.

For a comprehensive analysis of oocyte numbers at P16 and P21, we used NOBOX as an oocyte marker and employed a two-step approach. First, we manually generated a surface on the medulla region of the ovary, applied a mask excluding the cortex region, and established a new channel specifically for the analysis of growing oocytes in the medulla, utilizing an object size filter of 12 µm at P16 and 45 µm at P21. Second, we isolated only the cortex region of the whole ovary by setting the voxel values inside the surface to 0, created a new channel only for the cortex region and counted the number of NOBOX+ primordial and primary oocytes utilizing an object size filter of 4 µm at P16 and 6 µm at P21. GFP-labeled cells at P21 and at subsequent stages were meticulously chosen by visually inspecting each z-stack of whole-mount stained ovaries.

The quantification of oocytes in ovaries from 2, 6, and 12-month-old females was conducted using distinct oocyte markers, including VASA and NOBOX, as well as AMH, a marker for granulosa cells of growing follicles. We utilized object size filters to count growing follicles in adult and aged ovaries within the range of XY diameters from 15 µm to 45 µm. Oocytes in 2 month-old ovaries were counted by two independent scorers.

The analysis of LINE-1 ORF1p expression was conducted by selecting three representative z-stacks from each ovary. This approach was used due to segmentation limitations in Imaris with cytoplasmic expression in our automated whole-mount analysis pipeline.

### Fluorescence Activated Cell Sorting (FACS) of fetal oocytes

E17.5 fetal ovaries from pregnant *Rosa26^mT/mG^* dams (crossed with *Sycp3^CreERT^*^2^*^/+^*males) were dissected in ice cold 0.4% BSA in PBS, and each pair of ovaries was digested in 150 µl of 0.25% trypsin-EDTA (Fisher Scientific cat# 25200056) in 1.5-ml Eppendorf tubes at +37°C for 20 min. 15 min into the digestion, the suspension was gently triturated to mechanically dissociate the ovaries. After trypsin-EDTA incubation, the suspension was gently triturated and DNase I (1 mg/ml) was added at a 1:10 dilution, followed by additional pipette trituration. To quench the reaction, an equal volume of FBS (Gibco cat#10437028) was added to the cell suspension; Sytox Blue viability dye (Invitrogen cat#S34857) was added to samples at 1:1000 dilution, and then samples were filtered through a 35 μm filter into FACS tubes (Falcon cat#352235).

To isolate GFP+ and RFP+ cells from E17.5 *Sycp3^CreERT^*^2^*^/+^; Rosa26^mTmG/+^*ovaries, we utilized a BD FACSAria II system. Cells were gated on FSC height vs area comparison to remove doublets/multiplets, and then dead cells were gated on Sytox Blue positivity. GFP^high^, GFP^low^, and RFP+ (GFP^negative^) cells were sorted into 1.5-ml Eppendorf tubes containing 300 µl of 10% FBS in PBS. The total number of ovaries utilized and the numbers of sorted cells are depicted in **Extended Data Fig.2b**. After sorting, the samples were promptly processed to prepare meiotic spreads.

### Preparation, immunofluorescence staining, and analysis of chromosomal spreads

An equal volume of the hypotonic buffer [30 mM tris (pH 8.2), 50 mM sucrose, 17 mM sodium citrate, 5 mM EDTA, 0.5 mM dithiothreitol (DTT), and 0.5 mM phenylmethanesulfonyl fluoride] and sorted cells in solution were mixed then incubated for 30 min at room temperature. The suspension was centrifuged for 10 min at 1000 rpm. The supernatant was removed from each tube and the cells were resuspended in 100 mM sucrose. A hydrophobic barrier was drawn on positively charged glass microscope slides (precleaned in 70% EtOH) then equal volumes of fixative solution [1% PFA, 0.15% Triton X-100, and 3 mM DTT (pH 9.2)] and cell suspension were pipetted into the bordered area from approximately one foot above to make chromosome spreads. Slides were allowed to air-dry at room temperature under a laminar flow hood and then submerged in 0.4% Kodak Photo-flo 200 (Kodak Professional, cat #1464510) in Milli-Q H2O for 2 min twice. Last, slides were allowed to air-dry under the laminar flow hood and stored at −80°C until staining.

Slides were thawed then washed for 2 min with 0.4% Photo-flo 200 in Milli-Q H2O solution. Subsequently, slides were washed in 0.4% Photo-flo 200 in PBS then 0.1% Triton X-100 in PBS for 10 min each. Slides were blocked for 10 min in sterile filtered antibody dilution buffer (ADB) consisting of 10 ml normal goat serum, 3 g BSA, 50 μl Triton X-100, and 990 ml 1X PBS. Primary antibodies; MLH1, RAD51, SYCP1, SYCP3, and HORMAD1 (**Supplementary Table 2**) were diluted in ADB then 90 μl of primary antibody solution was applied to the chromosome spreads on the slide, covered with a parafilm coverslip, and incubated in a room temperature humid chamber overnight. Following incubation, slides were washed for 10 min in each of the following: 0.4% Photo-flo 200 in PBS, 0.1% Triton X-100 in PBS, and ADB. 90 μl of secondary antibody solution was applied to the chromosome spreads on the slide, covered with a parafilm coverslip, and incubated in a dark, room temperature humid chamber for 2 hours. Slides were washed three times for 5 min in 0.4% Photo-flo 200 in PBS then one time for 5 min in 0.4% Photo-flo 200 in Milli-Q H2O. 30 μl of Prolong Diamond antifade with DAPI was applied then covered with a glass coverslips. Slides were stored at 4°C prior to analysis. All images were captured on a Zeiss Axio Imager epifluorescence microscope at 63x magnification using standardized exposure times for each antibody condition and processed using Zeiss Zen Blue (version 3.0). Images were adjusted in ImageJ to standardize background across all images.

MLH1 and RAD51 foci that co-localized with SYCP3 in pachytene oocytes were counted by two independent scorers. Scores from cells with minor discrepancies were averaged; in the event of major MLH1 score discrepancies (more the 2 foci difference between scorers), the cell was excluded from analysis. Synapsis defects were categorized into four groups: complete synapsis (where all SYCP1 and SYCP3 co-localized without any fragmentation); partial and complete asynapsis (where part or all of the SYCP3 axial element did not co-localize with SYCP1); and fragmented (where SYCP1 and SYCP3 completely co-localize, but the SCs appear choppy or discontinuous).

### Breeding studies and analysis of fluorescence in progeny

For breeding studies, *Sycp3^CreERT^*^2^*^/+^; Rosa26^nTnG/+^* female mice that received a dose of 4-OHT at E9.5 by mouth pipetting were set up at 7 weeks of age with CD1 WT males. The females (total of 14 females across two cohorts) were monitored daily for births and maintained in continuous breeding with the males until they reached 12 months of age. The pups generated by continuous breeding were screened at P1-P3 with a SL10S spot lamp (Clare Chemical Research, Dark Reader Spot Lamp). The SL10S contains 3 high power blue LEDs and a proprietary blue filter. The peak wavelength of the SL10 is around 470 nm. The number of GFP+, RFP+, GFP-; RFP-pups were recorded.

### MII oocyte collection and RNA isolation

MII oocytes were collected from 3-week-old *Sycp3^CreERT^*^2^*^/+^; Rosa26^nTnG/+^* females that were superovulated by injection of 5 IU of pregnant mare serum gonadotropin (PMSG; Ilex Life Sciences, cat#A22721K) followed 48 hours later by 5 IU of human chorionic gonadotropin (hCG; Ilex Life Sciences, cat#A225005). 13 hours after hCG injection, females were euthanized, cumulus-oocyte complexes (COCs) were isolated from the oviduct in M2 media containing 4mg/ml BSA (Sigma cat#M7167), then COCs were incubated in M2 media containing 0.3 mg/ml Hyaluronidase (Sigma cat#H4272) for 1 min to remove cumulus cells. Denuded oocytes were washed in M2 media containing BSA and transferred to glass-bottom petri dishes (Thermo Scientific, cat#150680) containing a drop of M2 media containing BSA. GFP and RFP expression were checked under an Olympus IX71 microscope with an attached Lumencor Sola Light Engine. Oocytes were hand-picked based on their GFP expression and divided into 3 groups; GFP high, GFP low, and RFP. The following oocyte numbers were used for the 3 replicates for each group: GFP high: 15, 15, 16; GFP low: 49, 52, 57, and RFP: 88, 87, 100. RNA was extracted from oocytes using PicoPure RNA isolation kit (Thermo Fisher Scientific, Arcturus Pico Pure RNA Isolation Kit cat# KIT0204) according to the manufacturer’s instructions.

### RNA isolation and Bulk RNA seq

All sample QC, library preparation, sequencing, and data analysis were performed by Novogene as briefly described below.

Sample QC: RNA was quantified using Qubit RNA HS assay (Thermo Fisher, cat# Q32851), and RIN scores were calculated using Bioanalyzer 2100 Eukaryote Total RNA Nano (Agilent Technologies). Library preparation: Messenger RNA was purified from total RNA using poly-dT magnetic beads, and mRNA was fragmented. First strand cDNA synthesis was carried out using random hexamer priming, followed by second strand cDNA synthesis. Fragment ends were repaired, A-tailed, and ligated with sequencing adapters, followed by size selection, amplification via PCR, and purification. Final libraries were quantified using Qubit, and run on an Agilent Bioanalyzer system to check for proper size distribution. Post QC, index-barcoded libraries were pooled and paired-end sequencing was performed on an Illumina sequencing platform.

Data analysis: All data analysis was carried out by Novogene. Raw fastq reads were processed using the program fastp to remove adapter sequences, poly-N reads, and low quality reads (based on Q20, Q30, and %GC scores). Resulting cleaned reads were mapped to the mm10 reference genome (indexed using Hisat2) using the splice aware aligner Hisat2 (v2.0.5). The number of reads mapping to each gene was calculated using featureCounts v.1.5.0-p3. FPKM values were calculated for each gene, normalizing for gene length and sequencing depth. Differential expression analysis was performed using DESeq2 (v.1.20.0), and adjusted p-values were calculated using the Benjamini and Hochberg’s approach for controlling the false discovery rate. Genes with an adjusted p-value ≤ 0.05 were classified as differentially expressed.

### Immunofluorescence staining of MII oocytes

GFP+ and RFP+ oocytes were fixed with 4% PFA in PBS at +37°C for 20 min, washed three times with 0.1% Tween-20 in PBS (PBST) for 10 mins each, and permeabilized with 0.5% Triton X-100 in PBS for 20 min at room temperature. After washing with PBST three times for 10 mins each, oocytes were blocked with 3% BSA in PBST at +4°C for 4 hours and incubated with primary antibodies (**Supplementary Table 2**) diluted in blocking buffer [2% donkey serum, 0.1% BSA, 0.01% Tween-20 in PBS] at +4°C overnight. The next day, oocytes were washed 3 times with PBST for 10 min each at room temperature and incubated with Alexa Fluor–conjugated secondary antibodies in blocking buffer at room temperature for 2.5 hours. Oocytes were washed three times with PBST for 10 min each, transferred to glass-bottom dishes, and imaged using a white-light Leica TCS SP8 inverted confocal microscope with an HC PL APO CS2 63X/1.40 oil objective, 2X optic zoom, and 2 µm z step size.

### Statistical Analysis

Sample size used for experiments were based on prior experience with mouse embryonic, postnatal, adult, and aged ovary experiments that showed significance. Student’s t-test was used for analyzing spatial (anterior-posterior and medial-lateral) distributions of germ cells, percentage of GFP oocytes in primordial/primary and growing follicles, expression of different markers (cPARP and LINE-1 ORF1-p) between GFP+ vs GFP-cells. Differences in MLH1 and RAD51 counts among GFP^negative^g, GFP^low^, and GFP^high^ groups were analyzed by one-way ANOVA with a Tukey’s post-hoc test for multiple comparisons.The relative frequencies of Synapsis defects were analyzed among GFP^negative^-neg, GFP -^low^, and GFP-^high^ oocytes using a Fisher’s Exact test.

### Data and Code Availability

All data needed to evaluate the conclusions in the paper are present in the paper and/or the Supplementary Materials. Bulk RNA-seq data have been deposited in the Gene Expression Omnibus (GEO) under the accession code GSE272099. Code for analysis of fetal ovaries is available at https://github.com/BIDCatUCSF/Angular-Radial-Position-Distributi

## Extended Data Figure Legends

**Extended Data Fig. 1.**
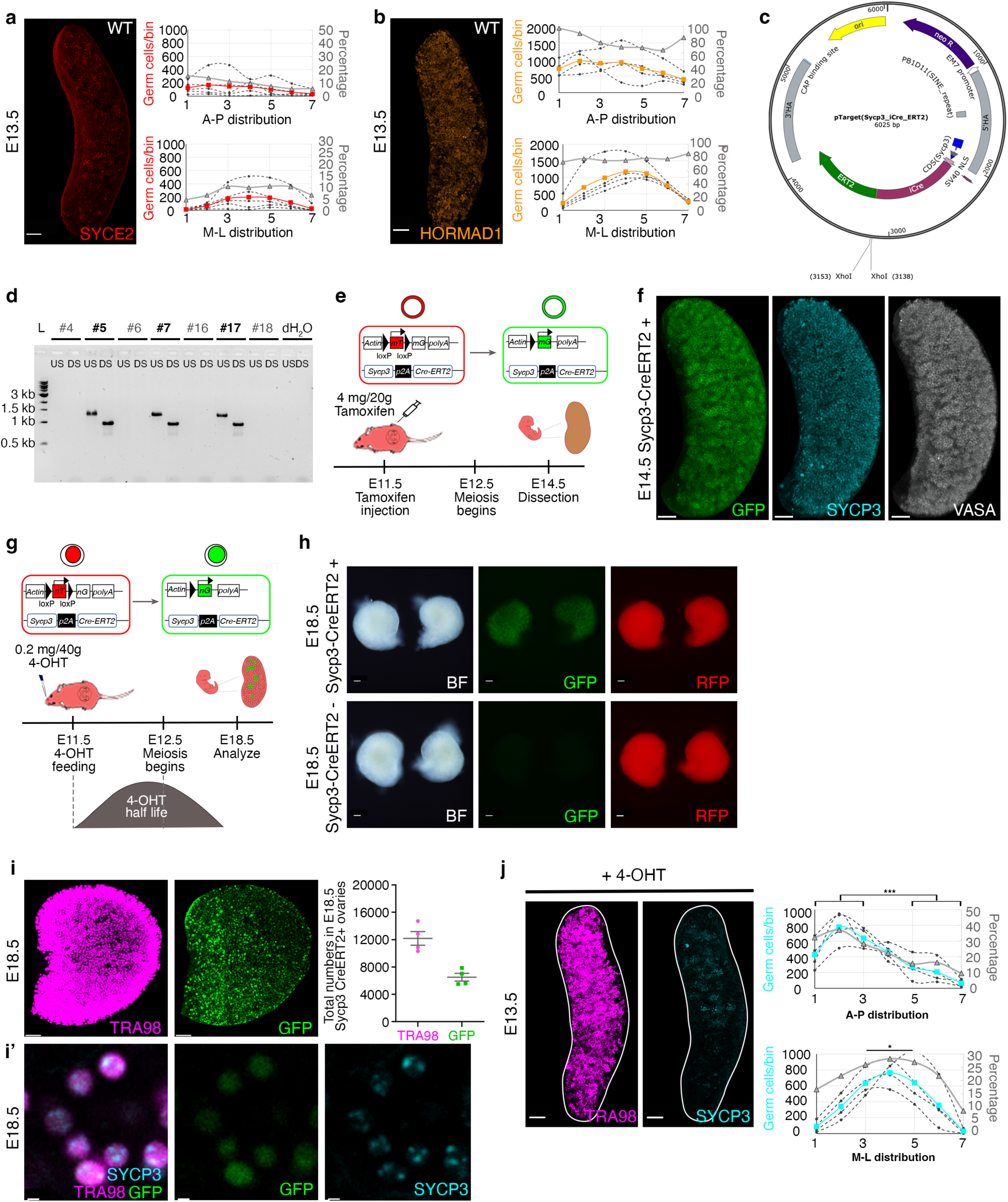
Spatial dynamics of additional markers of meiotic progression and validation of the Sycp3-CreERT2 lineage tracing system. (**a**) SYCE2 and (**b**) HORMAD1 expression exhibit no spatial bias in wholemount immunostained E13.5 wild-type ovaries. Scale bars, 50 µm. **c,** The targeting vector that was co-injected with CAS9 and sgRNA into FVB/N zygotes to generate the *Sycp3-CreERT2* line. **d,** Genotyping of founder mice positive for targeting (#5, #7, #17 in bold). L; ladder, US; Upstream junction, DS; Downstream junction. **e,** Illustration of the tamoxifen dosing regimen designed to evaluate the specificity and efficiency of the *Sycp3-CreERT2* line. Upon intraperitoneal tamoxifen injection at E11.5, the activated CreERT2 recombinase excises membrane-anchored tdTomato (mT) in Sycp3-expressing cells and initiates EGFP (mG) expression in cells, which leads to constitutive expression of membrane GFP. **f,** Whole-mount immunofluorescence detection of membrane GFP expression, observed in nearly all SYCP3 and VASA expressing oocytes at E14.5. Scale bars, 50 µm. **g,** Schematic for low dose oral administration of 4-OHT at E11.5, one day before meiotic initiation, which resulted in robust labeling in *Sycp3^CreERT^*^2^*^/+^; Rosa26^nT-nG/+^* ovaries at E18.5, shown at (**h**) low magnification, scale bars; 10 µm (**i**) as GFP was detected in over 50% of TRA98+ oocytes. Scale bars, 50 µm. (**i’**) GFP+ cells also expressed endogenous SYCP3 protein, confirming their meiotic status. Scale bars, 5 µm. **j,** 4-OHT administration at E11.5 had no effect on the wave of radial meiotic initiation as evaluated by endogenous SYCP3 protein expression analysis at E13.5. SYCP3 positive cells were predominantly located in the anterior (bins 1 to 3 on x axis of A-P distribution graph; \*\*\**P*=0.0001) and middle (bins 3 to 5 on x axis of M-L distribution graph; *P=0.0358) of E13.5 *Sycp3^CreERT^*^2^*^/+^; Rosa26^nT-nG/+^* ovaries (n=4). Scale bars, 50 µm.

**Extended Data Fig. 2.**
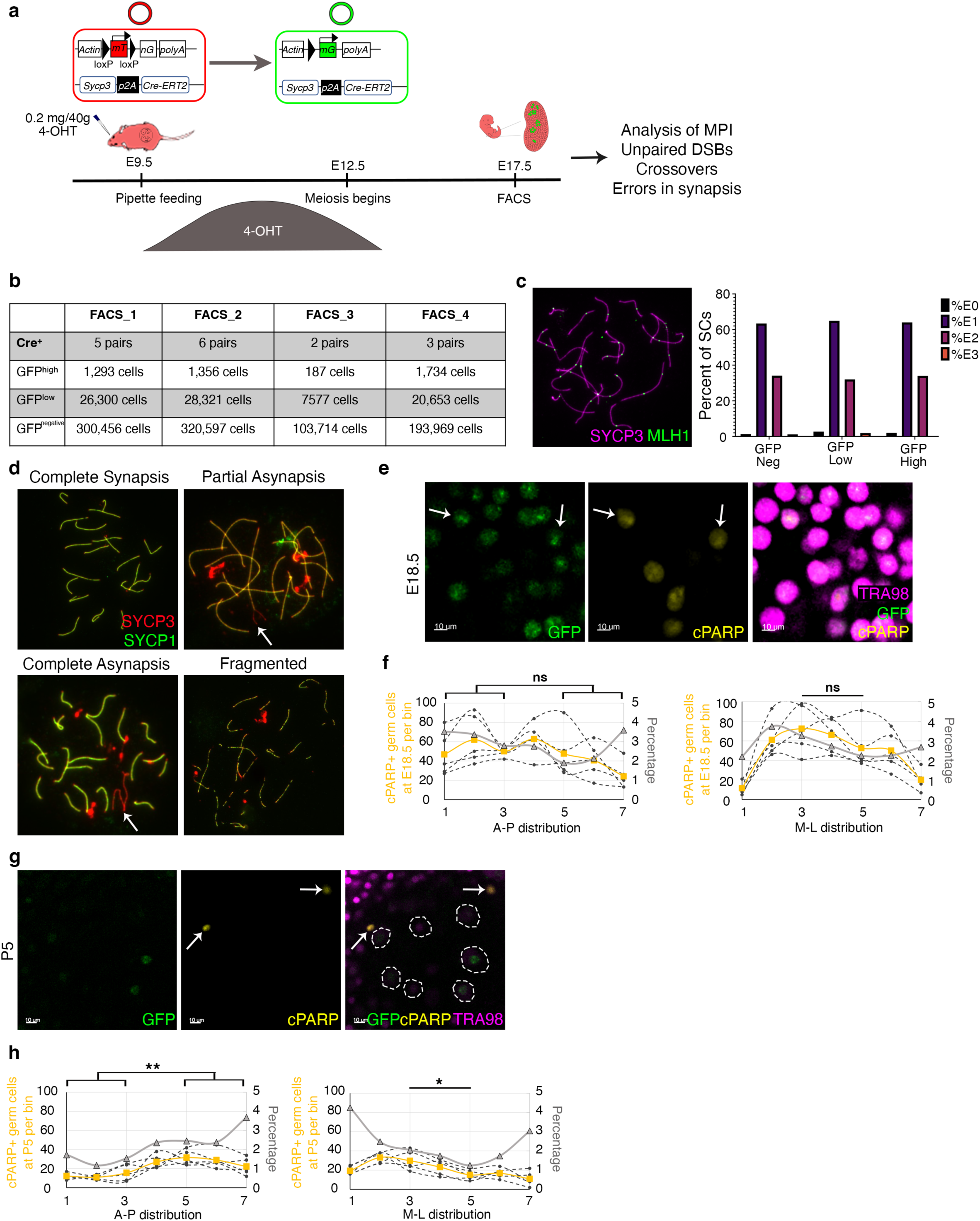
Chromosomal analysis and spatiotemporal dynamics of apoptosis in Sycp3-CreERT2 labeled oocytes. **a**, Experimental design for labeling the first meiotic entrants with membrane GFP, followed by downstream chromosomal analysis and (**b**) numbers of cells sorted for each (GFP^high^, GFP^low^, and GFP^negative^) group. Note that only GFP+ gates contain pure oocytes, while oocytes in GFP^negative^ gate were identified in meiotic spreads. **c,** Immunofluorescence labeling of SYCP3 and MLH1 and the frequency distribution of MLH1 foci across each synaptonemal complex (SC) within single nuclei. SCs were classified based on the number of MLH1 foci: those without an MLH1 focus were termed “E0” (0 exchanges), with one focus “E1” (1 exchange), with two foci “E2” (2 exchanges), and with three foci “E3” (3 exchanges). For E0, Chi square = 3.74, p = 0.15; for E1, Chi square = 0.44, p = 0.8; for E2, Chi square = 1.88, p = 0.39; and for E3, Chi square = 2.37, p = 0.31 as determined by Fisher’s Exact Test. GFP^negative^ oocytes exhibited a slightly lower frequency of E0s by comparison with GFP^low^ and GFP^high^ oocytes (1.3%, 2.7%, and 2.0%, respectively), though this difference was not statistically significant (Chi square = 3.74; p = 0.15). **d,** Pachytene oocytes were stained for SYCP3 and SYCP1 to assess the synapsis quality: complete synapsis (SYCP1 and SYCP3 perfectly co-localize without errors), partial asynapsis (white arrow pointing to a forked region of asynapsis), complete asynapsis (white arrow pointing to two axial elements without any SYCP1), and fragmented SC (SYCP1 and SYCP3 co-localize, but the SCs appear broken). **e,** Section view from whole-mount imaging of E18.5 *Sycp3^CreERT^*^2^*^/+^; Rosa26^nT-nG/+^* ovaries, white arrows show GFP and cPARP co-expressing TRA98+ oocytes. Scale bars, 10 µm. **f,** cPARP+ oocytes were randomly distributed in E18.5 *Sycp3^CreERT^*^2^*^/+^; Rosa26^nT-nG/+^* ovaries. **g,** Section view from whole-mount imaging of P5 *Sycp3^CreERT^*^2^*^/+^; Rosa26^nT-nG/+^* ovaries. Dashed white circles delineate growing follicles. cPARP expression is concentrated within primordial follicles (arrows) at P5. Scale bars, 10 µm. **h,** A higher proportion of TRA98+ oocytes in the posterior (\*\**p*=0.0019) and lateral regions (cortex; \**p*=0.0261) of P5 ovaries exhibited cPARP expression.

**Extended Data Fig. 3.**
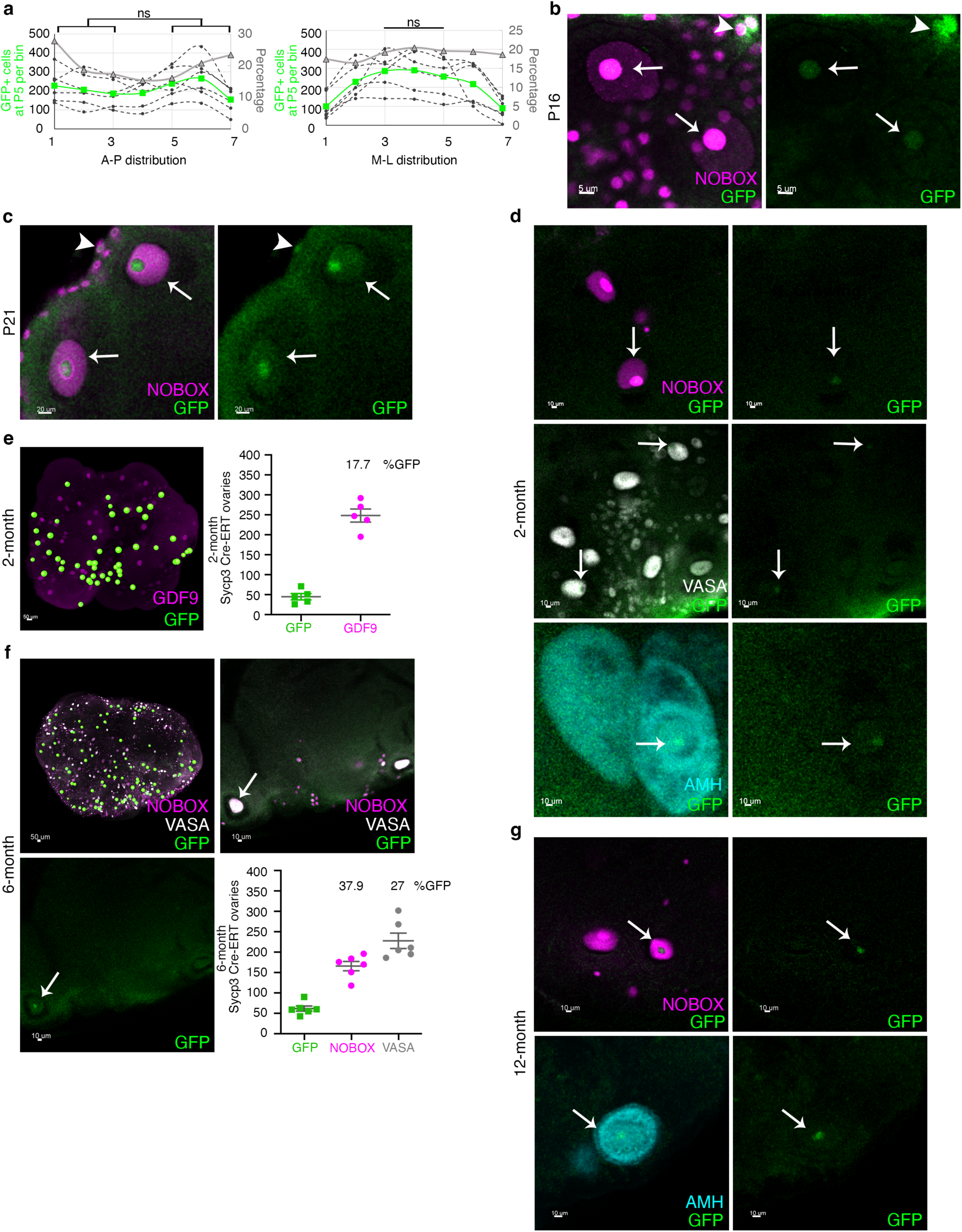
Localization studies of Sycp3-CreERT2 lineage traced oocytes in postnatal and adult ovaries. **a**, Distribution of GFP+ oocytes in *Sycp3^CreERT^*^2^*^/+^; Rosa26^nT-nG/+^* ovaries at P5. (**b)** and **(c**) High magnification images showing GFP+ growing follicles (arrows) and GFP+ primordial/primary follicles (arrowheads) in P16 and P21 whole-mount stained ovaries from *Sycp3^CreERT^*^2^*^/+^; Rosa26^nT-nG/+^* mice. Oocytes in prepubertal ovaries were labeled with a NOBOX antibody. Scale bars are 5 µm for P16 and 20 µm for P21. **d,** High magnification images of NOBOX (top, magenta) , VASA (middle, gray), and AMH (bottom, cyan) expressing GFP+ growing follicles (arrows) in *Sycp3^CreERT^*^2^*^/+^; Rosa26^nT-nG/+^* ovaries at 2-month. Scale bars, 10 µm. **e,** Detection and quantification of GDF9 and GFP expression at 2-month (n=5). Scale bar, 50 µm. **f,** Whole-mount immunofluorescence and quantification of NOBOX, VASA, and GFP expressing growing follicles in *Sycp3^CreERT^*^2^*^/+^; Rosa26^nT-nG/+^* ovaries (n=6) at 6-month. Scale bars, 50 µm. **g,** High magnification images showing GFP+ growing follicles (arrows) in 12-month-old ovaries expressing NOBOX (top, magenta) and AMH (bottom, cyan).

**Extended Data Fig. 4.**
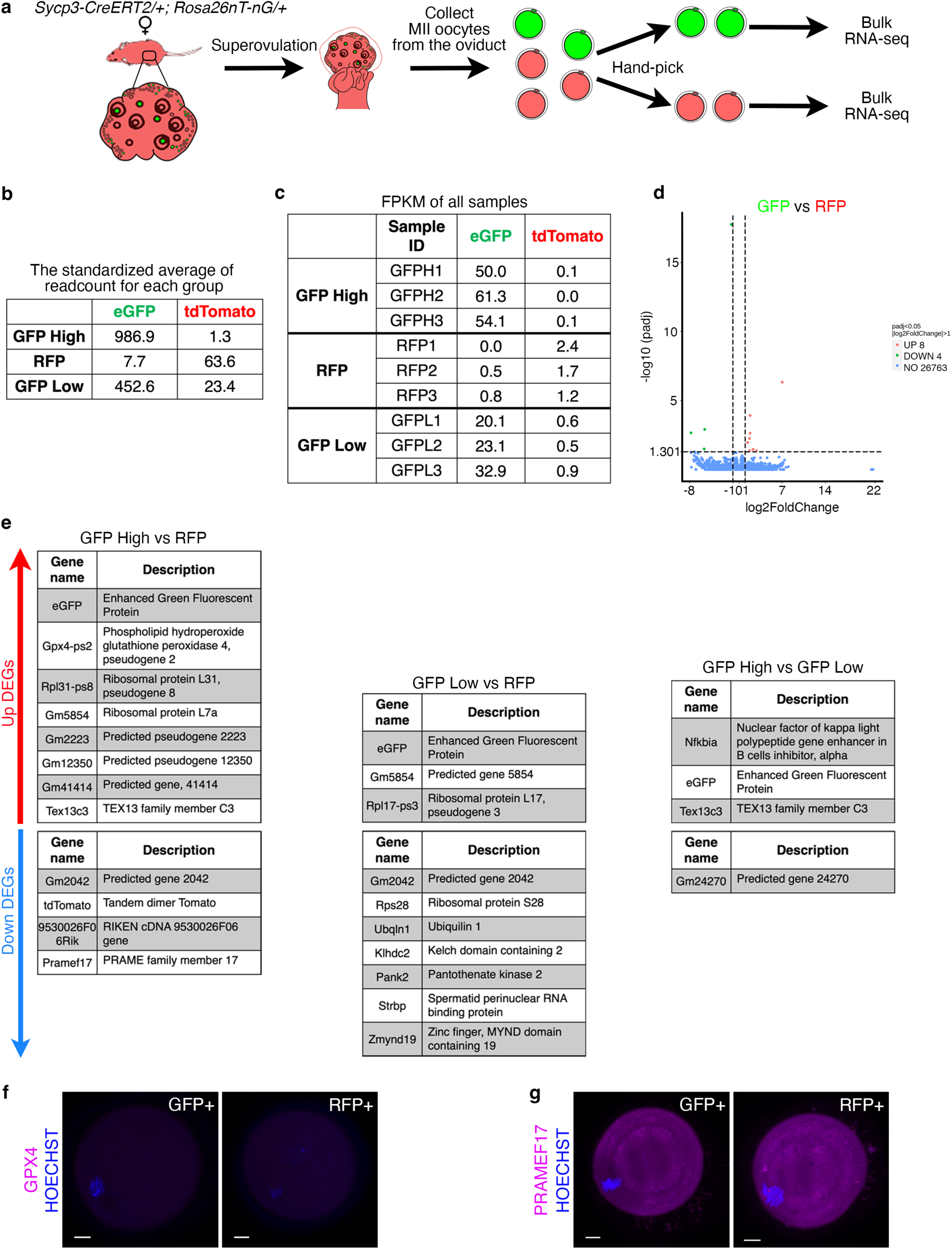
Transcriptional signature of mature Sycp3-CreERT2 lineage traced oocytes collected at P21 is unremarkable. **a**, Experimental approach to isolate GFP+ and RFP+ (GFP-) MII oocytes for Bulk RNA-seq analysis. *Sycp3^CreERT^*^2^*^/+^; Rosa26^nT-nG/+^* females exposed to 4-OHT *in utero* were superovulated at P21 and MII oocytes were manually selected for transcriptome analysis. Two fluorescence intensities were detected in MII GFP+ oocytes: GFP high and GFP low. Oocytes with low GFP signal were kept as a separate group, and transcriptomic analysis was carried out on the three groups (GFP high, GFP low, and RFP+). Further analysis was primarily focused on the GFP high and RFP+ groups. **b,** The standardized average of readcount for each group. **c,** Fragments Per Kilobase per Million mapped fragments (FPKM) for each sample. **d,** A volcano plot showing a very small number of differentially expressed genes (8 upregulated and 4 downregulated) between GFP high and RFP+ oocytes (adjusted p-value ≤ 0.05). **e,** Differentially expressed genes between GFP high vs RFP, GFP low vs RFP, and GFP high vs GFP low. Immunofluorescence labeling of (**f**) GPX4 and (**g**) PRAMEF17 proteins in MII oocytes revealed similar expression in both GFP high and RFP+ oocytes. Scale bars, 10 µm.

**Supplementary Table 1.** Quantification of number of oocytes at each time point.

**Supplementary Table 2.** Primers and Primary antibodies used in the study.

**Supplementary Movie 1. Identification of GFP+ growing follicles in the postnatal ovary.** Slice view of the P5 ovary shows that oocytes in primordial follicles are marked by TRA98 (magenta). GFP+ growing follicles (indicated by arrows) are distinguished by decreased TRA98 expression and the presence of AMH (cyan) secreted by the surrounding granulosa cells.

**Supplementary Movie 2. Detection of GFP+ oocytes in the intact prepubertal ovary.** 3D projection and slice view of NOBOX+ oocytes (magenta) provide a comprehensive analysis of the total oocyte population within the intact P16 ovary. The spot view, merged with the immunofluorescence signal, allows for the easier detection of representative GFP+ oocytes within growing follicles (arrows) and primordial/primary follicles (arrowheads).

**Supplementary Movie 3. Identification of GFP+ oocytes in the prepubertal ovary.** Slice view of the P21 ovary reveals nuclear GFP+ oocytes in growing follicles co-labeled with NOBOX (arrows). Conversely, very few GFP+ oocytes are observed in primordial/primary follicles, which are marked by TRA98 (arrowheads).

**Supplementary Movie 4. GFP+ growing follicles in the adult ovary.** Slice view of the 2-month-old ovary shows AMH+ small growing follicles containing nuclear GFP+ oocytes (arrows).

## Acknowledgments

This work was funded by NIH grants 1DP2OD007420 (DJL), 1R01GM122902 (DJL), 1R01ES023297 (DJL), 1F31HD110208 (MHF), 1F31HD108875 (EAG), F31HD101234 (SAC), R01HD041012 (PEC), K99112986 (TSH) as well as the Global Consortium for Reproductive Health through the Bia-Echo Foundation GCRLE-0123 (DJL) and GCRLE-1620 (BS), and the W.M. Keck Foundation (DJL). We thank all members of the Laird lab for their support and input, Dr. Amy Laird for statistical advice, the UCSF Parnassus FlowCytometry Core for assistance with cell sorting, and Dr. Aleks Rajkovic for kindly gifting the NOBOX antibody.

## Author contributions

Conceptualization: DJL, BS, Methodology: BS, EAG, MHF, SC, AW, TH, Funding acquisition: DJL, PC, Supervision: DJL, Writing – original draft: DJL, BS, Writing – review & editing: DJL, BS, EAG, MHF, PC

### Competing interests

DJL is a Scientific Advisor for Vitra, Inc.

