## Supplementary material for "Sustained fertility from first-wave follicle oocytes that pause their growth": Table 1

| E16.5 | | | | | |
|---|---|---|---|---|---|
| **In Figure** | **Litter** | **Ovary ID** | **GFP** | **TRA98** | **%GFP** |
| 1 | 1 | 1 | 301 | 18,155 | 1.7 |
| | | 2 | 1090 | 19,848 | 5.5 |
| | | 3 | 817 | 15,642 | 5.2 |
| | | 4 | 655 | 18,884 | 3.5 |
| | | 5 | 1597 | 19,366 | 8.2 |
| | | 6 | 1059 | 18,756 | 5.6 |
| | 2 | 1 | 683 | 11,268 | 6.1 |
| | | 2 | 1077 | 13,236 | 8.1 |
| | 3 | 1 | 1021 | 14,604 | 7.0 |
| | | 2 | 829 | 12,596 | 6.6 |
| | | **Mean** | **912.9** | **16,236** | **5.8** |

| E18.5 | | | | | |
|---|---|---|---|---|---|
| | **Litter** | **Ovary ID** | **GFP** | **TRA98** | **%GFP** |
| 1 | 1 | 1 | 1371 | 12,358 | 11.09 |
| | | 2 | 1102 | 12,389 | 8.89 |
| | 2 | 1 | 1445 | 10,002 | 14.45 |
| | | 2 | 1350 | 10,446 | 12.92 |
| | | 3 | 1388 | 10,776 | 12.88 |
| | 3 | 1 | 1955 | 16479 | 11.86 |
| | | 2 | 1590 | 14368 | 11.07 |
| | | **Mean** | **1457.3** | **12,403** | **11.9** |

| P5 | | | | | | | | | | | |
|---|---|---|---|---|---|---|---|---|---|---|---|
| | **Litter** | **Ovary ID** | **GFP cortex** | **GFP growing follicles** | **Total GFP** | **TRA98 cortex** | **TRA98 growing follicles** | **Total TRA98** | **%GFP cortex** | **%GFP growing follicles** | **%GFP (Overall)** |
| 3, Extended Data 3 | 1 | 1 | 946 | 66 | 1,012 | 3,733 | 444 | 4,177 | 25.3 | 14.9 | 24.2 |
| | | 2 | 753 | 29 | 782 | 7,112 | 198 | 7,310 | 10.6 | 14.6 | 10.7 |
| | | 3 | 1,691 | 63 | 1,754 | 9,555 | 229 | 9,784 | 17.7 | 27.5 | 17.9 |
| | | 4 | 1,800 | 80 | 1,880 | 7,741 | 247 | 7,988 | 23.3 | 32.4 | 23.5 |
| | | 5 | 1,590 | 67 | 1,657 | 9,934 | 176 | 10,110 | 16.0 | 38.1 | 16.4 |
| | 2 | 1 | 1,772 | 50 | 1,822 | 8,177 | 205 | 8,382 | 21.7 | 24.4 | 21.7 |
| | | **Mean** | **1,425** | **59.2** | **1,485** | **7,709** | **249.8** | **7,959** | **19.1** | **25.3** | **19.1** |

| P16 | | | | | | | | | | | |
|---|---|---|---|---|---|---|---|---|---|---|---|
| | **Litter** | **Ovary ID** | **GFP cortex** | **GFP growing follicles** | **Total GFP** | **NOBOX cortex** | **NOBOX growing follicles** | **Total NOBOX** | **%GFP cortex** | **%GFP growing follicles** | **%GFP (Overall)** |
| 3, Extended Data 3 | 1 | 1 | 88 | 96 | 184 | 9,090 | 301 | 9,391 | 1.0 | 31.9 | 2.0 |
| | | 2 | 182 | 100 | 282 | 8,587 | 252 | 8,839 | 2.1 | 39.7 | 3.2 |
| | 2 | 1 | 138 | 47 | 185 | 4,814 | 248 | 5,062 | 2.9 | 19.0 | 3.7 |
| | | 2 | 210 | 71 | 281 | 6,167 | 295 | 6,462 | 3.4 | 24.1 | 4.3 |
| | | 3 | 105 | 70 | 175 | 6,218 | 245 | 6,463 | 1.7 | 28.6 | 2.7 |
| | | 4 | 245 | 69 | 314 | 5,934 | 246 | 6,180 | 4.1 | 28.0 | 5.1 |
| | | **Mean** | **161.3** | **75.5** | **237** | **6,802** | **264.5** | **7,066** | **2.5** | **28.5** | **3.5** |

| P21 | | | | | | | | | | | |
|---|---|---|---|---|---|---|---|---|---|---|---|
| | **Litter** | **Ovary ID** | **GFP cortex** | **GFP growing follicles** | **Total GFP** | **NOBOX cortex** | **NOBOX growing follicles** | **Total NOBOX** | **%GFP cortex** | **%GFP growing follicles** | **%GFP (Overall)** |
| | 1 | **1** | 10 | 52 | 62 | 5126 | 315 | 5441 | 0.2 | 16.5 | 1.1 |
| | | **1** | 20 | 70 | 90 | 6,528 | 277 | 6,805 | 0.3 | 25.3 | 1.3 |

| | | | | | | | | | | | |
|---|---|---|---|---|---|---|---|---|---|---|---|
| 3, Extended Data 3 | 2 | **2** | 13 | 64 | 77 | 4,115 | 247 | 4,362 | 0.3 | 25.9 | 1.8 |
| | | **3** | 7 | 36 | 43 | 4,582 | 256 | 4,838 | 0.2 | 14.1 | 0.9 |
| | 3 | **1** | 6 | 99 | 105 | 5,338 | 275 | 5,613 | 0.1 | 36.0 | 1.9 |
| | | **2** | 3 | 111 | 114 | 5,820 | 244 | 6,064 | 0.1 | 45.5 | 1.9 |
| | | **Mean** | **9.8** | **72.0** | **81.8** | **5,252** | **269.0** | **5,521** | **0.2** | **27.2** | **1.5** |

**2-month**

| | | **Ovary ID** | **GFP** | **NOBOX** | **%GFP** |
|---|---|---|---|---|---|
| 3, Extended Data 3 | 1 | 1 | 103 | 307 | 33.6 |
| | 2 | 1 | 40 | 248 | 16.1 |
| | | 2 | 57 | 268 | 21.3 |
| | 3 | 1 | 56 | 328 | 17.1 |
| | | 2 | 50 | 281 | 17.8 |
| | | 3 | 67 | 272 | 24.6 |
| | | 4 | 45 | 337 | 13.4 |
| | 4 | 5 | 60 | 239 | 25.1 |
| | | **Mean** | **59.8** | **285** | **21.1** |

| | | **Ovary ID** | **GFP** | **VASA** | **AMH** | **%GFP (based on VASA)** | **%GFP (based on AMH)** |
|---|---|---|---|---|---|---|---|
| 3, Extended Data 3 | 1 | 1 | 67 | 341 | 316 | 19.6 | 21.2 |
| | | 2 | 42 | 313 | 303 | 13.4 | 13.9 |
| | | 3 | 62 | 319 | 320 | 19.4 | 19.4 |
| | | 4 | 37 | 226 | 176 | 16.4 | 21.0 |
| | | 5 | 31 | 255 | 189 | 12.2 | 16.4 |
| | | **Mean** | **47.8** | **291** | **260.8** | **16.2** | **18.4** |

| | **Litter** | **Ovary ID** | **GFP** | **GDF9** | **%GFP** |
|---|---|---|---|---|---|
| 3, Extended Data 3 | 1 | 1 | 26 | 195 | 13.3 |
| | | 2 | 33 | 270 | 12.2 |
| | | 3 | 71 | 292 | 24.3 |
| | | 4 | 51 | 247 | 20.6 |
| | | 5 | 43 | 237 | 18.1 |
| | | **Mean** | **44.8** | **248** | **17.7** |

**6-month**

| | | **Ovary ID** | **GFP** | **VASA** | **NOBOX** | **%GFP (based on VASA)** | **%GFP (based on NOBOX)** |
|---|---|---|---|---|---|---|---|
| 3, Extended Data 3 | 1 | 1 | 90 | 268 | 184 | 33.6 | 48.9 |
| | | 2 | 63 | 302 | 196 | 20.9 | 32.1 |
| | 2 | 1 | 43 | 205 | 170 | 21.0 | 25.3 |
| | | 2 | 55 | 211 | 176 | 26.1 | 31.3 |
| | | 3 | 59 | 195 | 118 | 30.3 | 50.0 |
| | | 4 | 60 | 186 | 151 | 32.3 | 39.7 |
| | | **Mean** | **61.7** | **227.8** | **165.8** | **27.3** | **37.9** |

**12-month**

| | Litter | **Ovary ID** | **GFP** | **NOBOX** | **AMH** | **VASA** | **%GFP (based on NOBOX)** | **%GFP (based on AMH)** | **%GFP (based on VASA)** |
|---|---|---|---|---|---|---|---|---|---|
| 3, Extended | 1 | 1 | 13 | 72 | 54 | | 18.06 | 24.07 | |
| | | 2 | 19 | 98 | 66 | | 19.39 | 28.79 | |
| | | 3 | 15 | 84 | 73 | | 17.86 | 20.55 | |

| | | | | | | | | | |
|---|---|---|---|---|---|---|---|---|---|
| Data 3 | 2 | 1 | 16 | 96 | | 85 | 16.67 | | 18.82 |
| | 3 | 1 | 11 | 93 | | 120 | 11.83 | | 9.17 |
| | | 2 | 11 | 57 | | 69 | 19.30 | | 15.94 |
| | 4 | 3 | 15 | | 88 | 89 | | 17.05 | 16.85 |
| | | 4 | 5 | | 22 | 32 | | 22.73 | 15.63 |
| | | 5 | 12 | | 63 | 72 | | 19.05 | 16.67 |
| | | **Mean** | **13.0** | **83.3** | **61.0** | **77.8** | **17.2** | **22.0** | **15.5** |
