## Supplementary material for "Sustained fertility from first-wave follicle oocytes that pause their growth": Table 2

<table>
<tr><th colspan="4">PRIMERS</th></tr>
<tr><td>Sycp3CreERT2 Upstream (US) Forward</td><td colspan="2">5'-TGT TCC ATT AGA TAG CTA CAG GAG T-3'</td><td rowspan="2">Expected size:<br>1273 bp</td></tr>
<tr><td>Sycp3CreERT2 Upstream (US) Reverse</td><td colspan="2">5'-CTT CGA CAT CCC CTG CTT GT -3'</td></tr>
<tr><td>Sycp3CreERT2 Downstream (DS) Forward</td><td colspan="2">5'-CAT CTA TAC TTG GAA CCT TGT ACC TTG-3'</td><td rowspan="2">Expected size:<br>956 bp</td></tr>
<tr><td>Sycp3CreERT2 Downstream (DS) Reverse</td><td colspan="2">5'-AGA TCC TGA TGA TTG GTC TCG TCT G-3'</td></tr>
<tr><td>nTnG Wild Type</td><td colspan="2">5'-GGA GCG GGA GAA ATG GAT ATG-3'</td><td rowspan="3">Expected sizes:<br>Mutant: 320 bp<br>WT: 603 bp</td></tr>
<tr><td>nTnG Mutant</td><td colspan="2">5'-CCA GGC GGG CCA TTT ACC GTA AG-3’</td></tr>
<tr><td>nTnG Common</td><td colspan="2">5'-AAA GTC GCT CTG AGT TGT TAT-3’</td></tr>
<tr><td>mTmG Wild Type Forward</td><td colspan="2">5’-AGG GAG CTG CAG TGG AGT AG-3’</td><td rowspan="3">Expected sizes:<br>Mutant: 128 bp<br>WT: 212 bp</td></tr>
<tr><td>mTmG Mutant Forward</td><td colspan="2">5’-TAG AGC TTG CGG AAC CCT TC-3’</td></tr>
<tr><td>mTmG Common</td><td colspan="2">5’-CTT TAA GCC TGC CCA GAA GA-3’</td></tr>
<tr><th colspan="4">PRIMARY ANTIBODIES</th></tr>
<tr><th>Protein</th><th>Host Species</th><th>Company #Catalogue number</th><th>Dilution</th></tr>
<tr><td>GFP</td><td>Chicken</td><td>Aves #1020</td><td>1:200</td></tr>
<tr><td>TRA98</td><td>Rat</td><td>Abcam #ab82527</td><td>1:200</td></tr>
<tr><td>SYCP3</td><td>Mouse</td><td>Abcam #ab97672</td><td>1:200</td></tr>
<tr><td>SYCP3</td><td>Rabbit</td><td>Cohen Lab at Cornell*</td><td>1:10,000</td></tr>
<tr><td>SYCP1</td><td>Rabbit</td><td>Abcam #ab15087</td><td>1:1000</td></tr>
<tr><td>SYCE2</td><td>Rabbit</td><td>Invitrogen #PA5-59857</td><td>1:200</td></tr>
<tr><td>HORMAD1</td><td>Rabbit</td><td>Proteintech #13917-1-AP</td><td>1:200</td></tr>
<tr><td>RAD51</td><td>Rabbit</td><td>EMD Millipore #PC-130</td><td>1:500</td></tr>
</table>

| | | | |
|---|---|---|---|
| MLH1 | Mouse | BD Biosciences 550838 | 1:50 |
| cPARP | Mouse | BD Pharmingen #558710 | 1:100 |
| LINE-1 ORF1p | Rabbit | Abcam #ab216324 | 1:200 |
| AMH | Goat | Santa Cruz #sc-6886 | 1:500 |
| AMH | Rabbit | Abcam #ab272221 | 1:200 |
| VASA | Rabbit | Abcam #ab13840 | 1:200 |
| NOBOX | Goat | Rajkovic lab at UCSF | 1:1000 |
| GDF9 | Goat | ThermoFisher Scientific #PA5-47924 | 1:200 |
| GPX4 | Rabbit | Abcam #ab125066 | 1:200 |
| PRAMEF17 | Rabbit | Invitrogen #PA5-98082 | 1:200 |

*Lenzi, M. L. et al. Extreme heterogeneity in the molecular events leading to the establishment of chiasmata during meiosis i in human oocytes. Am. J. Hum. Genet. 76, 112–127 (2005).
